# Human Adenovirus (HAdV) mediated inhibition of the tumor suppressor p53 is controlled by SUMOylation of the viral E2A/DBP protein

**DOI:** 10.1101/2025.02.07.636424

**Authors:** Miona Stubbe, Hans Christian Stubbe, Julia Mai, Verena Plank, Samuel Hofmann, Lilian Göttig, Thomas Günther, Adam Grundhoff, Stefan Krebs, Peter Groitl, Julia Mayerle, Ron T. Hay, Thomas Dobner, Sabrina Schreiner

**Affiliations:** Institute of Virology, School of Medicine, Technical University of Munich, Germany; Institute of Virology, Medical Center – University of Freiburg, Freiburg, Germany; Medical Depertment II, University Hospital, LMU, Munich, Germany; German Center for Infection Research, DZIF, partner site Munich, Germany; Institute of Virology, Hannover Medical School, Hannover, Germany; Cluster of Excellence RESIST (Resolving Infection Susceptibility; EXC 2155), Hannover Medical School, Hannover, Germany; Division of Pediatric Neurology and Metabolic Medicine, Center for Pediatric and Adolescent Medicine, University Hospital Heidelberg, Heidelberg, Germany; Leibniz Institute of Virology, LIV, Hamburg, Germany; Laboratory for Functional Genome Analysis, Gene Center, LMU Munich, Germany; Wellcome Trust Centre for Gene Regulation and Expression, College of Life Sciences, University of Dundee, Dundee, United Kingdom

**Keywords:** HAdV, E2A/DBP, p53, SUMOylation, PML-NBs, tumor suppressor

## Abstract

HAdV express early viral genes to modulate the activity of the cellular tumor suppressor p53 and ensure efficient viral replication, through various processes, whilst some of them are not completely understood. HAdV oncoprotein E1B-55K interactions with SUMO, PML-IV/V and Sp100A are tightly connected to the transforming potential of the viral factor in non-lytic infections. SUMO modified HAdV DNA binding protein (DBP) E2A interacts with PML and Sp100A acting as a molecular bridge between viral replication centers (RCs) and PML tracks. Here, we provide a novel concept showing that HAdV E2A is involved in the inhibition of p53 activity and apoptosis; thus we identified a novel route of exhibiting virus induced oncogenic potential. We revealed an E1B-55K independent localization of p53 within HAdV replication centers marked by E2A. In the absence of any functional E1B-55K protein expression, p53 interacts with E2A. E2A SUMOylation promotes HAdV- mediated inhibition of p53/DNA binding and p53-dependent transactivation by supporting p53 degradation, E1B-55K interaction and colocalization with PML-V that leads to E1B-55K induced upregulation of p53 SUMOylation. In sum, we provide evidence for a novel mechanism by which the HAdV DBP E2A protein inhibits p53- mediated transcriptional activation and apoptotic host cell response. Intriguingly, these findings change the longstanding dogma of host DNA damage response (DDR) inactivation by HAdV infection and offer crucial insights to improve future p53 selective oncolytic virus therapies and adenoviral vectors.

## INTRODUCTION

p53 is a cellular tumor suppressor with a key role in maintenance of genome integrity during oncogenic processes and ongoing DNA virus replication (1). Activation of the p53 protein induces cell cycle arrest or apoptosis is primarily triggered as a response to various stress signals, such as DNA damage and oncogene activation (2–4). Even though under basal conditions p53 degradation limits its functions, p53 still binds to specific target genes and controls their activity (5, 6). p53 was originally discovered as a target of SV40 polyomavirus large T-antigen (7–10), but due to the role of p53 in antiviral innate immunity (11), many other DNA viruses also developed elaborate mechanisms to counteract the activity of this host factor (reviewed in 12, 13-20). Despite decades of research, exact modes of viral modulation of p53 pathway are still not fully understood.

Adenoviruses are classified as DNA tumor viruses due to their oncogenic properties in non-permissive mammalian cells (21). The DNA of several Human Adenovirus (HAdV) types has been found in different human tumors (22, 23), but solid causative association between HAdV infection and oncogenicity in humans has not been established yet. However, it has been recently reported that HAdV-C5 E1A/E1B oncogenes lead to a complete transformation of human mesenchymal stem/stroma cells (hMSCs) in tissue culture (24). HAdV infection is usually mild and self-limiting in immunocompetent individuals, however it can lead to severe outcomes in immunocompromised patients (25–27). Additionally, infection with the recently identified HAdV 14p1 type can be lethal even in healthy patients (28). Specific treatment against HAdV infection is not available, but promising progress in this direction has been made by recent discoveries showing strong antiviral capacity of arsenic trioxide in cell culture models by promoting PML-NB integrity (29). On the other hand, HAdV are thoroughly researched in context of their therapeutic potential. AdV vectors are the most used vectors in the field of gene therapy (30, 31) and oncolytic AdV are identified as a novel class of anticancer agents (32, 33). Furthermore, AdV are extensively investigated and used as vaccine vectors (34, 35) against pathogens such as Ebola (36), HIV (37) or currently emerging SARS-CoV-2 (38–40). Nevertheless, the understanding of the complex virus-host interplay during HAdV infection is far from understood in detail especially for various serotypes others than HAdV-C5.

HAdV employ different approaches to manipulate the p53 activity and establish productive infection. HAdV early protein E1A leads cells from G1- to S-phase to provide optimal conditions for viral replication (19, 41–44). This causes p53 stabilization (19, 45–48), which would induce apoptosis without the action of HAdV oncoproteins such as E1B-55K, E4orf6 and E4orf3, that all inhibit p53 using distinct processes (19, 49, 50). HAdV early protein E1B-55K inhibits p53 function via several different mechanisms. E1B-55K directly interacts with the tumor suppressor, inhibiting p53-dependent transcription (51–53). The E1B-55K nuclear export signal (NES) and the SUMO conjugation motif (SCM) are essential for nuclear-cytoplasmic relocalization of p53 mediated by E1B-55K that completely abrogates the p53 activity (54, 55). It has been reported that p53 can be found together with E1B-55K in sites resembling to HAdV replication centers (RCs) (56), but direct connection between p53 and the HAdV RCs has so far not been fully investigated. Moreover, E1B-55K prevents PCAF (p300/CBP-associated factor) - mediated p53 acetylation necessary for high DNA binding affinity via interaction with the two cellular proteins (57). E1B-55K interaction with PML-IV and PML-V leads to p53 SUMOylation and inhibition of p53 transcriptional activity (58, 59). Furthermore, SUMO modified E1B-55K specifically binds, SUMOylates and sequesters Sp100A isoform into the insoluble nuclear matrix or into cytoplasmic inclusions. When E1B-55K/Sp100A complex recruits p53, Sp100A stimulated p53 transcriptional activity is effectively eliminated (60). Additionally, E1B-55K associates with E4orf6 to assemble the E3 ubiquitin ligase complex, which among other antiviral factors also degrades p53 (61–63).

The early HAdV DBP E2A, promotes diverse functions during HAdV life cycle, such as viral gene expression (64), mRNA stability (65), virion assembly (66) and maturation (67). Due to its essential role in DNA replication, E2A is termed the marker protein for HAdV RCs (68). HAdV RCs localize juxtaposed to PML-NBs reorganized into tracks by the early viral protein E4orf3 (69–75). E2A SUMOylation is a molecular mechanism connecting HAdV RCs and PML tracks, used by the virus to exploit beneficial host components within PML tracks and to promote viral replication (76, 77). E1B-55K functions in inhibition of p53-mediated transcription, resulting in transformation of cells in cooperation with E1A, are tightly linked to E1B-55K SUMO PTM and interactions with PML proteins and Sp100A (55, 58–60). Similarly to E1B-55K, E2A also interacts with PML proteins and Sp100A. E2A SUMOylation-dependent binding to these cellular factors leads to positioning of PML tracks in proximity to viral RCs (77). However, a clear association between E2A and HAdV- mediated p53 inactivation has so far not been established.

Here we demonstrate a novel mechanism used by HAdV to inhibit p53 activity through the action of the HAdV DNA-binding protein E2A. Our results revealed p53 localization within HAdV RCs, independent on E2A SUMOylation and E1B-55K. In the absence of functional E1B-55K, E2A interacts with p53. E2A SUMOylation even regulates HAdV-mediated inhibition of p53-dependent transactivation and apoptosis by promoting p53 degradation and E1B-55K interaction with PML-V necessary for p53 SUMOylation with subsequent inhibition of the tumor suppressor transcription activating properties.

## MATERIAL AND METHODS

### Cell culture

A549 (78), HepaRG (79; Thermo Scientific), HepaRG cells stably expressing His/HA-SUMO-2 (kindly provided by Prof. Roger Everett/University of Glasgow), HEK-293 (Sigma-Aldrich Inc) and 2E2 (80) cells were grown in DMEM medium supplemented with 10% FCS or 5% FCS in case of A549 cells, 100 U of penicillin and 100 µg of streptomycin per ml at 37°C in atmosphere with 5% CO_2_. HepaRG cells were propagated in media additionally supplemented with 5 μg/ml of bovine insulin and 0.5 μM of hydrocortisone. HepaRG-SUMO-2-His/HA cells were maintained under 2 µM puromycin selection. 2E2 cells were maintained in media supplemented with 90 µg/µl hygromycin B and 250 µg/µl of G-418. To induce helper function media used for propagation of 2E2 cells was additionally supplemented with 1 µg/ml of doxycycline. All cell lines were consistently tested for mycoplasma contamination.

### Plasmids, mutagenesis, and transient transfection

N–terminally EYFP-tagged human PML-V isoform construct was kindly provided by Prof. Roger Everett/University of Glasgow. For transient transfections, a mixture of DNA and 25 kDa linear polyethylenimine (*Polysciences*) was applied to subconfluent cells as previously described (81).

### Viruses

In this work, H5*pg*4100 was used as the wild type HAdV-C5 parental virus (HAdV wt) (82). HAdV E2A SCM mutant virus was generated as it was previously reported in (83). This virus contains point mutations in the E2A gene that exchange amino acids at positions 94, 132 and 202 from lysine to arginine, leading to disruption of the SCM in E2A (77). H5*pm*4149 virus that contains four stop codons in the E1B-55K region (AS position 3, 8, 86 and 88) was used as HAdV E1B-55K depleted (E1B-55K minus) virus (84). H5*pm*4154, containing a stop codon in the E4orf6 region (AS 66) was used as HAdV E4orf6 depleted (E4orf6 minus) virus (61). H5*pm*4101, HAdV-C5 E1B-55K mutant with three amino acids exchanges (L83/87/91A) within the NES of the E1B-55K sequence was used as HAdV E1B-55K NES virus (84). H5*pm*4102, HAdV-C5 E1B-55K mutant containing one amino acid exchange from lysine to arginine (K104R) within the SCM of the E1B-55K sequence was used as HAdV E1B-55K SCM virus (84). Viruses were propagated in H1299 and HEK 293 cells and titrated in 2E2 (83) and HEK 293 cells as previously described (80).

### Protein analysis and antibodies (Ab)

Cells were harvested according to the experimental setup and resuspended in RIPA buffer freshly supplemented with 0.2 mM PMSF, 1 mg/ml Pepstatin A, 5 mg/ml aprotinin, 20 mg/ml leupeptin, 25 mM iodacetamide and 25 mM N-ethylmaleimide as previously described (85). The cell lysates were incubated 30 min on ice, sonicated for 30 s, and the insoluble debris was pelleted at 11,000 rpm/4°C. For immunoprecipitation, samples were precleared by adding Pansorbin cells (*Millipore Calbiochem*) and Protein A sepharose beads (1 mg/sample). Protein A sepharose beads (3 mg/sample) were coupled with 0.25 μl of monoclonal mouse antibody raised against p53 protein (anti-p53 DO-1) or with 0.25 µl of polyclonal rabbit anti-GFP Ab, for 1 h at 4°C. Protein A sepharose beads coupled with desired antibody were added to precleared samples and rotated for 1 h at 4°C. Proteins attached to the antibody-coupled protein A sepharose were centrifuged, washed three times with RIPA buffer, boiled for 5 min at 95°C in 2x Laemmli buffer and subsequently analysed by Western blotting as previously reported (86). To detect p53 SUMO PTM, five 100 mm tissue culture dishes (*Sarstedt*) of HepaRG cells stably expressing His/HA-SUMO-2 were treated according to the experimental setup, pooled together and subjected to denaturing purification and analysis of SUMO conjugates performed as described in (86, 87). Primary Abs specific for HAdV proteins used in this study included E2A mouse mAb B6-8 (87), E1B-55K mouse mAb 2A6 (85) and E4orf6 rat mAb RSA3 (Kindly provided by HMGU monoclonal antibody unit). Primary Ab specific for cellular proteins included monoclonal mouse Ab against β-actin AC-15 (A5441; *Sigma-Aldrich*), monoclonal mouse antibody raised against p53 (DO-1; *Santa Cruz Biotechnology*) and polyclonal rabbit antibody raised against p53 (FL-393, *Santa Cruz Biotechnology*). Polyclonal rabbit Ab against anti-GFP/YFP (ab290; *Abcam*) and monoclonal mouse Ab against the 6xHis epitope (631213; *Clontech*) were used to label proteins carrying specific tags. Secondary Ab conjugated to HRP (*horseradish peroxidase*) used to detect proteins by immunoblotting were anti-rabbit IgG and anti-mouse IgG (*Jackson/Dianova*).

### Indirect immunofluorescence

Cells were cultivated on glass cover slips and transfected and/or infected. Immunofluorescence studies were performed using previously defined protocols (84). Cytoskeleton wash buffer (CSK buffer) at pH 6.8, containing 100mM NaCl, 300 mM Sucrose, 3 mM MgCl_2_ * 6H2O and 10 mM PIPES, supplemented with TritonX-100 to final concentration of 0.5%, was used to remove soluble cytoplasmic proteins and loosely held nuclear proteins. Digital images were acquired using a *Nikon TiE* microscope equipped with *Perkin Elmer UltraView Vox System*. *Pearson’s correlation* coefficient, determined with *Volocity 6.2.1* software (*Perkin Elmer*), was used to analyse colocalization of E2A and p53 and E1B-55K and YFP-PML-V. Images were cropped and assembled with *Affinity Publisher 2*.

### Chromatin immunoprecipitation (ChIP)

24h after infection of HepaRG cells with HAdV wt or HAdV E2A SCM, chromatin was crosslinked by incubation with 1% formaldehyde. The reaction was stopped by the addition of glycine at a final concentration of 0.125 M. After 5 min of incubation, cells were washed once in ice-cold PBS. Protease inhibitors were added in all buffers in final concentrations of 1 μg/ml leupeptin, 1 μg/ml pepstatin,10 U/ml aprotinin, and 1 mM PMSF. Cross linked cells were rotated for 10 min at 4°C in Buffer I (50mM Hepes-KOH, 140 mM NaCl, 1 mM EDTA, 10% Glycerol, 0.5% Nonidet P-40 and 0.25% Triton X-100). After centrifugation at 2000g for 10 min at 4°C, pelleted nuclei were resuspended in 1ml Buffer II (10 mM Tris-HCl, 200mM NaCl, 1 mM EdTA and 0.5 mM EGTA) and rotated for 10 min at 4°C. For lysis nuclei were resuspended in Buffer III (1% SDS, 10 mM EDTA, 50 mM Tris-HCl, pH 8). Isolated chromatin was sheared by using a Bioruptor Pico (Diagenode) (15 cycles, in 30 seconds intervals on/off). 10% Triton X-100 was added and cell debris was pelleted by centrifugation at 20,000g, for 10 min at 4°C. Supernatant with isolated and chromatin was collected. For immunoprecipitation, chromatin of 1×10^6^ cells was diluted 1:10 in dilution buffer (0.01% SDS, 1.1% Triton X-100, 1.2 mM EDTA, 16.7 mM Tris-HCl, 167 mM NaCl) and 5 μg of the antibody specific for p53 (mAb DO-1, Santa Cruz) and the corresponding control mouse IgG antibody (Merck) were added and rotated for 20 h at 4°C. Antibody-chromatin complexes were precipitated by addition of 50 μl BSA-blocked magnetic Protein-A/G beads (Thermo Fisher) and incubated while rotating for 1 h at 4°C. Magnetic stand was used for washing steps. Washing of the beads was performed using 1 ml of the following salt buffers: low-salt buffer (0.1% SDS, 1% Triton X-100, 2 mM EDTA, 20mM Tris-HCl, 150 mM NaCl), high-salt buffer (0.1% SDS, 1% Triton X-100, 2 mM EDTA, 20 mM Tris-HCl, 500 mM NaCl) and LiCl-wash buffer (0.25 M LiCl, 1% Nonidet P-40, 1% Na-deoxycholate, 1 mM EDTA, 10 mM Tris-HCl). After washing the beads once with each salt buffer, TE buffer (10mM Tris-HCl, 1mM EDTA) was used for two subsequent washing steps. Chromatin was eluted by incubating beads in 210 μl elution buffer (50 mM Tris-HCl pH 8.0, 10 mM EDTA, 1% SDS) on a thermal block at 65°C and 1,000 rpm for 30 min. Magnetic stand was used to separate the beads from the chromatin-containing supernatant that was transferred to a reaction tube with 8 μl of 5 M NaCl. De-crosslinking was performed for both ChIP and input DNA samples diluted in 200 μl elution buffer, at 65°C and 1,000 rpm, overnight. Input DNA samples were processed equally as the ChIP samples starting with the de-crosslinking step. To degrade RNA, samples were incubated at 37°C for 2 h with added 200 μl TE buffer with 8 μl RNAse A (10 mg/ml, Sigma Aldrich). To degrade proteins, 7 μl of 300 mM CaCl_2_ and 4 μl proteinase K (20 mg/ml, Apli-Chem) were added to the samples and incubated for 1 h at 55°C. DNA purification was performed by phenol-chloroform-isoamyl alcohol (Roth) extraction in phase-lock gel tubes (5Prime, Quantabio) and precipitation with ethanol. ChIP DNA pellets and input DNA samples were diluted in 55 μl elution buffer. Obtained DNA was later subjected to ChIP-seq library preparation.

### ChIP-seq library preparation and sequencing

DNA samples (fragmented Chromatin after IP and input controls) were quantified by Qubit measurements. 10 µl of each sample was subjected to library generation via the NEB DNA Ultra II (product number: E7645L) protocol according to manufacturer’s instructions. Quality controlled libraries were sequenced with the NextSeq 1000/2000 P2 Reagents (100 Cycles) kit (product number: 20046811) in a paired-end mode (2x61 bp).

### ChIP-seq analysis

Paired end sequencing reads of all conditions were aligned to the human genome (hg38) and the Ad5 genome using bowtie (88). p53 enriched sites (p53 peaks) on the human genome were called in all samples using MACS2 (89) with standard settings and the respective input samples as controls. Peaks of all conditions were merged into a combined p53 peak set using bedtools (90). Combined peaks were annotated with the annotatePeaks.pl script of the HOMER sequencing analysis package (91). Differential p53 binding between conditions was detected with DiffReps (92) using presets of nucleosome size detection and G-test statistics. Differential sites were then overlapped with the annotated consensus peak set. Normalized heatmap visualization of all consensus peaks was generated with EaSeq (v1.1.1) (93). Motif detection within the consensus peakset was performed using the MEME-ChIP (94).

### Illumina RNA sequencing

Total RNA was isolated from whole-cell lysates with TRIzol reagent (*Invitrogen*) as described by the manufacturer. Strand specific, polyA-enriched RNA sequencing was performed as previously reported (95). RNA integrity number (RIN) was determined with the *Agilent 2100 BioAnalyzer* (*RNA 6000 Nano Kit*, Agilent). For library preparation, 1 μg of RNA was poly(A) selected, fragmented, and reverse transcribed using the *Elute, Prime, Fragment Mix* (Illumina). A-tailing, adaptor ligation, and library enrichment were performed according to the *TruSeq Stranded mRNA Sample Prep Guide* (Illumina). Quality and quantity of RNA libraries were evaluated using the *Agilent 2100 BioAnalyzer* and the *Quant-iT PicoGreen dsDNA Assay Kit* (Life Technologies). RNA libraries were sequenced as 150 bp paired end runs on an *Illumina HiSeq4000* platform at Helmholtz Zentrum München RNAseq Core Facility. Kallisto (version 0.46.1) was used to quantify transcript abundancies from paired-end RNA-seq reads (96). The index for Kallisto-pseudoalignment was built using the Genome Reference Consortium Human Build 38 patch release 13 (GRCh38.p13), (97). Differential gene expression was analyzed using the R package sleuth (version 1.46.0), (98).

### Caspase activity

The caspase activity was detected using the caspase-Glo® 3/7 assay kit (Promega) according to the manufacturer’s instructions. HepaRG and A549 cells were seeded into a white wall clear bottom 96-well plate at a density of 1.5 × 10^4^ cells/well in triplicate wells and cultured overnight. The cells were then ether uninfected or infected with HAdV wt or HAdV E2A SCM viruses for 48 h. HepaRG cells were infected with an MOI of 50 and A549 with an MOI of 20. At 48 h post infection HepaRG and A549 cells were treated with Actinomycin D (HepaRG 0.5µM, A549 10µM) in total 50µl suspension for 4.5h. 50 μL of caspase-Glo® reagent was added to each well afterward. The samples were incubated for 60 min at room temperature and the luminescence was measured using a Tecan (Männedorf, Switzerland) Infinite 200M plate reader.

### Statistical analysis

Testing for statistically significant differences in medians was performed using a two-sided *Mann-Whitney-U test* if not otherwise notified. All statistical analysis was carried out using the R language and environment for statistical computing, v4.2.1. Results were adjusted for multiple comparison using the Benjamini-Hochberg correction if applicable.

## RESULTS

### p53 localizes to viral RCs during HAdV infection

Subcellular localization is one of the main properties regulating the p53 functions (99–101). During HAdV infection, E1B-55K relocalizes p53 into E1B-55K containing sites that resemble to HAdV RCs (56). Nevertheless, the association between p53 and HAdV RCs has so far not been fully established. To investigate the influence of HAdV infection on p53 subcellular localization, we infected human liver and lung cells with HAdV wt virus and after 24 h performed immunofluorescence analysis (Fig. 1). Our data showed clear p53 colocalization with the HAdV RCs in infected HepaRG cells (Fig. 1 A). This observation was supported by the Pearsońs correlation coefficient for p53 and E2A, which was 0.71, above the border value for colocalization of 0.5 (Fig. 1 B). As expected based on previous reports (61–63), at 36 h post infection of HepaRG cells, it was not possible to detect p53 localization in RCs due to the onset of HAdV mediated p53 degradation (Suppl. Fig. 1). In infected A549 cells p53 was found in the region of HAdV RCs but it was more diffusely distributed when compared to the localization in HepaRG cells (Fig. 1 C). Nonetheless, Pearsońs correlation coefficient of 0.57 confirmed the colocalization between p53 and E2A (Fig. 1 D).

**Fig. 1.**
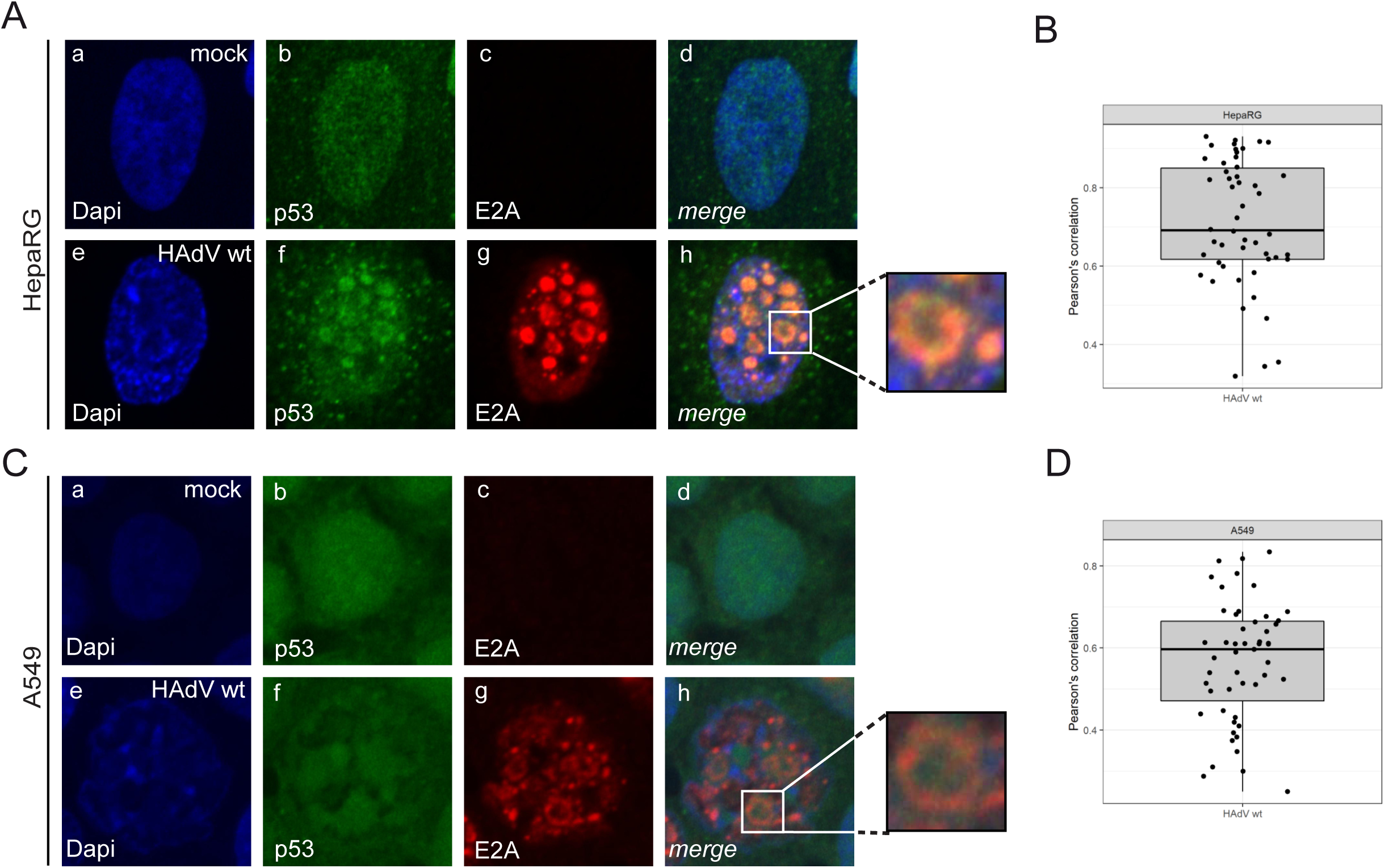
p53 colocalizes with the viral RCs during HAdV infection. (A, **C)** HepaRG **(A)** and A549 **(C)** cells were infected with HAdV wt virus at a multiplicity of infection of 20 FFU/cell. The cells were fixed with 2% PFA 24 h p.i. and double labeled with mAb rb (α-E2A) and mAb DO-1 (α-p53). Primary Abs were detected with Alexa647 (E2A, red) and Alexa488 (α-p53, green) conjugated secondary Abs. Nuclear staining was performed using Dapi. Anti-p53 (green; panels b, f) and anti-E2A (red; panels c, g), staining patterns representative of at least 50 analyzed cells are shown. Overlays of single images (merge) are shown in panels d and h. **(B, D)** Software Volocity was used to determine Pearson’s correlation coefficient for p53 and E2A in HepaRG **(B)** and A549 **(D)** cells infected with HAdV wt virus. For each cell, Pearson’s correlation was calculated for colocalization of p53 and E2A. The distributions of the respective correlation coefficients are displayed in box plots. Values above 0.5 are considered as colocalized.

### RC association of p53 is independent on E1B-55K

Throughout the course of HAdV infection, E1B-55K shuttles between cytoplasm and nucleus and localizes both in the nucleus and in perinuclear bodies (102). Inactivation of NES within E1B-55K leads to the complete redistribution of the viral protein from the cytoplasm to the nucleus, where it accumulates at the periphery of HAdV RCs. E1B-55K with a non-functional NES exhibits elevated levels of SUMO-1 posttranslational modification and increased rate of E1B-55K mediated transformation (54). Moreover, SUMOylation of E1B-55K is required for E1B-55K nuclear localization (84). The lack of E1B-55K SCM abolishes its capacity to transform cells in cooperation with E1A (55). E1B-55K mainly performs its oncogenic functions by modulation of the tumor suppressor p53 (51, 52, 58, 59). Our data show that p53 localizes in HAdV RCs during HAdV wt infection (Fig. 1) and it has been reported that E1B-55K changes p53 localization during HAdV infection (56). Therefore, we monitored the influence of E1B-55K on p53 localization in HAdV RCs by intracellular immunofluorescence studies. As E2A represents the marker protein for HAdV RCs (68) and it is modified by SUMO (77), we also investigated the effect of E2A SUMOylation on p53 localization during infection. We infected cells with HAdV wt, HAdV E2A SCM, HAdV E1B-55K minus, HAdV E1B-55K NES and E1B-55K SCM viruses. Our results obtained in immunofluorescence studies substantiated the previous findings mentioned above, showing the E1B-55K shuttling between cytoplasm and nucleus during HAdV wt infection ((102), Fig. 2 A, panels h and j, Fig.2 C). E2A SCM mutations did not affect the E1B-55K localization in nucleus and perinuclear bodies ((102), Fig. 2 A, panels m and o, Fig. 2 C). In concordance with previous reports, the inactivation of E1B-55K NES caused the accumulation of the viral protein in the HAdV RCs and PML tracks ((54), Fig. 2, panels w and y, Fig. 2 C), and the E1B-55K SCM mutation abrogated E1B-55K nuclear localization ((84), Fig. 2 A, panels b1 and d1, Fig. 2 C). Intriguingly, during HAdV wt infection we found p53 in HAdV RCs as well as in perinuclear bodies together with E1B-55K, independently on E2A SUMO PTM (Fig. 2 A, panels g, j and l, o, Fig. 2 C). Infection with the HAdV E1B-55K minus virus led to entirely nuclear p53 localization, with 68% of cells containing p53 in the HAdV RCs (Fig. 2 A, panels q and t, Fig. 2 C), and 32% of cells with diffuse p53 distribution. During HAdV E1B-55K NES virus infection, p53 localized exclusively in nucleus within HAdV RCs and PML tracks (Fig. 2 A, panels v and y, Fig. 2 C). The E1B-55K SCM mutation supported p53 localization together with E1B-55K in perinuclear bodies, but in all infected cells p53 was also found in HAdV RCs (Fig. 2 A, panels a1 and d1, Fig. 2 C). Pearsońs correlation coefficients determined for p53 and E2A signals substantiated the data we obtained in immunofluorescence studies (Fig. 2 B). We did not detect significant difference between Pearsońs correlation coefficients for p53/E2A determined during infection with HAdV E2A SCM virus compared to HAdV wt infection. Although they still colocalized, Pearsońs correlation coefficients for p53/E2A were significantly lower during infection with HAdV E1B-55K minus, HAdV E1B-55K NES and E1B-55K SCM viruses then during HAdV wt infection (Fig. 2 B).

**Fig. 2.**
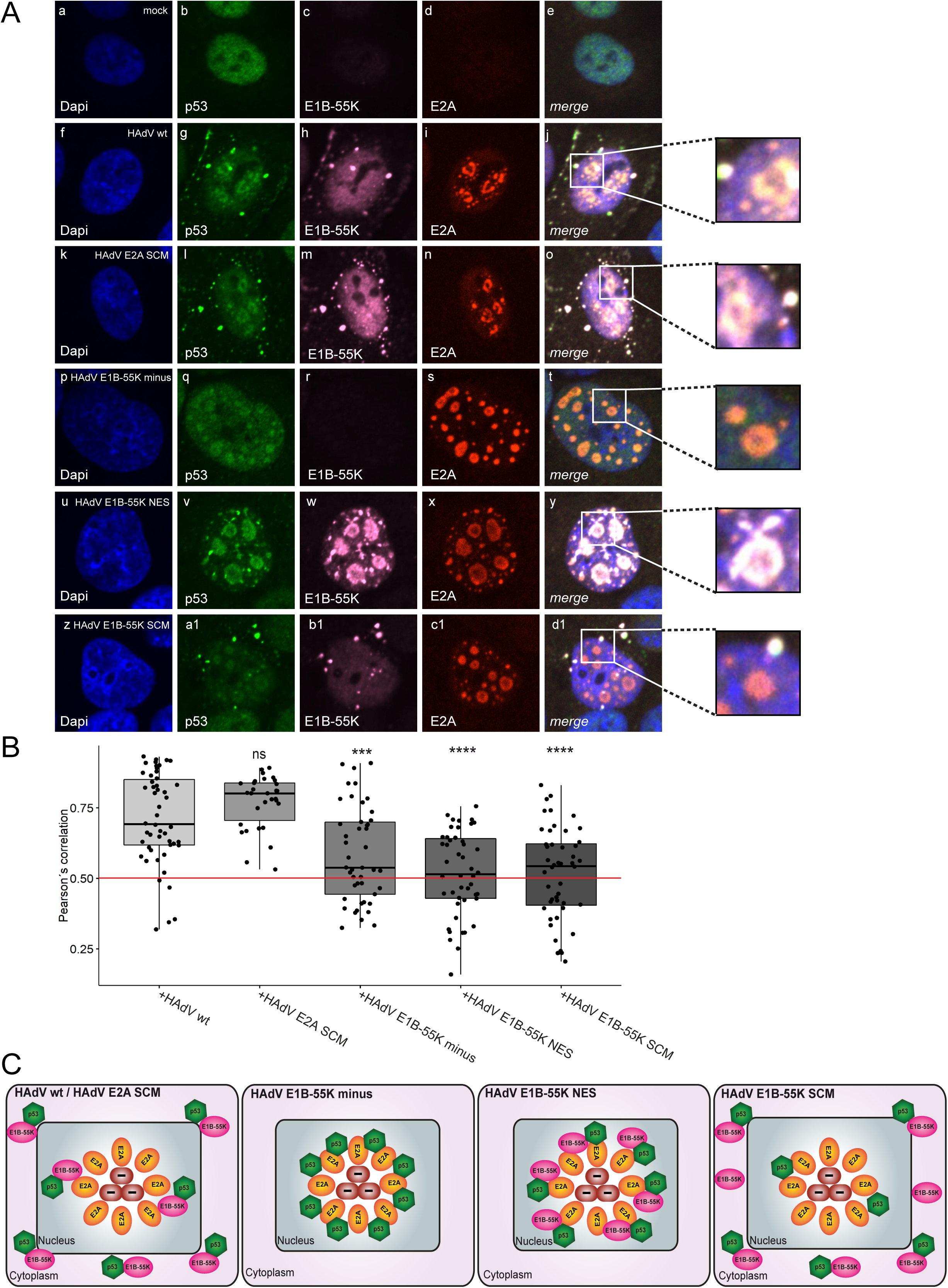
E1B-55K does not influence the p53 localization in HAdV RCs. **(A)** HepaRG cells were infected with HAdV wt, HAdV E2A SCM, HAdV E1B-55K minus, HAdV E1B-55K NES and E1B-55K SCM viruses at a multiplicity of infection of 20 FFU/cell. The cells were fixed with 2% PFA 24 h p.i. and triple labeled with mAb DO-1 m (α-p53), mAb 4E8 rat (α-E1B-55K) and mAb rb (α-E2A). Primary Abs were detected with Alexa488 (α-p53, green), Alexa640 (α-E1B-55K, pink) and Alexa568 (α-E2A, red) conjugated secondary Abs. Nuclear staining was performed using Dapi. Anti-p53 (green; panels b, g, l, q, v, a1), anti-E1B-55K (pink; panels c, h, m, r, w, b1) and anti-E2A (red; panels d, i, n, s, x, c1), staining patterns representative of 50 analyzed cells are shown. Overlays of single images (merge) are shown in panels e, j, o, t, y, d1. **(B)** Pearson’s correlation coefficient for E2A and p53 signals stained by mAb rb (α-E2A) and mAb DO-1 m (α-p53), in cells infected with HAdV wt, HAdV E2A SCM, HAdV E1B-55K minus, HAdV E1B-55K NES and E1B-55K SCM viruses was determined using the Volocity software. For each cell, Pearson’s correlation was calculated for colocalization of E2A with p53. The distributions of the respective correlation coefficients are displayed in a box plot. Values above 0.5 are considered colocalization. Statistical significances for differences in determined E2A/p53 Pearson’s correlation coefficient between HAdV wt infected cells and HAdV E2A SCM, HAdV E1B-55K minus, HAdV E1B-55K NES or E1B-55K SCM virus infected cells, were determined using two-sided Mann-Whitney-U test. ***, P < 0.001; ****, P < 0.0001; ns, not significant. **(C)** Schematic representation of p53 and E1B-55K localization during infection with HAdV wt, HAdV E2A SCM, HAdV 55K minus, HAdV E1B-55K NES and E1B-55K SCM viruses.

### E1B-55K prevents E2A binding to p53 in HAdV infected cells

Our data showed that p53 localizes in HAdV RCs independently on E1B-55K (Fig. 2). E2A is the marker protein for HAdV RCs (68) and a target of the host SUMOylation machinery (77). As SUMO PTM affects protein-protein interactions of its substrates (103) and E1B-55K influences p53 localization during infection (56), we investigated the interaction of E2A with p53 and its dependency on E2A SUMOylation and E1B-55K. We infected cells with HAdV wt, HAdV E2A SCM, HAdV E1B-55K minus, HAdV E1B-55K NES and E1B-55K SCM viruses. Immunoprecipitation analysis of p53 and subsequent staining for E2A did not show binding between the two investigated proteins during infection with HAdV wt or HAdV E2A SCM virus (Fig. 3 A, right panel, lanes 3 and 4, Fig. 3 B and C). However, E2A/p53 interaction was found in cells infected with HAdV E1B-55K minus virus (Fig. 3 A, right panel, lane 5, Fig. 3 B and C). Furthermore, the inactivation of E1B-55K NES allowed the examined interaction (Fig. 3 A, right panel, lane 6 Fig. 3 B and C). When E1B-55K SCM was impaired, E2A also interacted with p53 (Fig. 3 A, right panel, lane 7, Fig. 3 B and C). Densiometric analysis revealed similar amount of E2A pulled down with p53 antibody in cells infected with HAdV E1B-55K minus and HAdV E1B-55K NES viruses (Fig. 3 A, right panel, lanes 5 and 6, Fig. 3 B, Fig. 3 C). Although E2A was bound to p53 in HAdV E1B-55K SCM virus infected cells, the amount of pulled down E2A was reduced to 27% when compared to HAdV E1B-55K minus virus infected cells (Fig. 3 A, right panel, lane 7, Fig. 3 B). These results suggest that E1B-55K protein expression prevents E2A binding to p53 (Fig. 3 C), possibly to counteract p53 mediated disruption of E2A functions vital for viral replication.

**Fig. 3.**
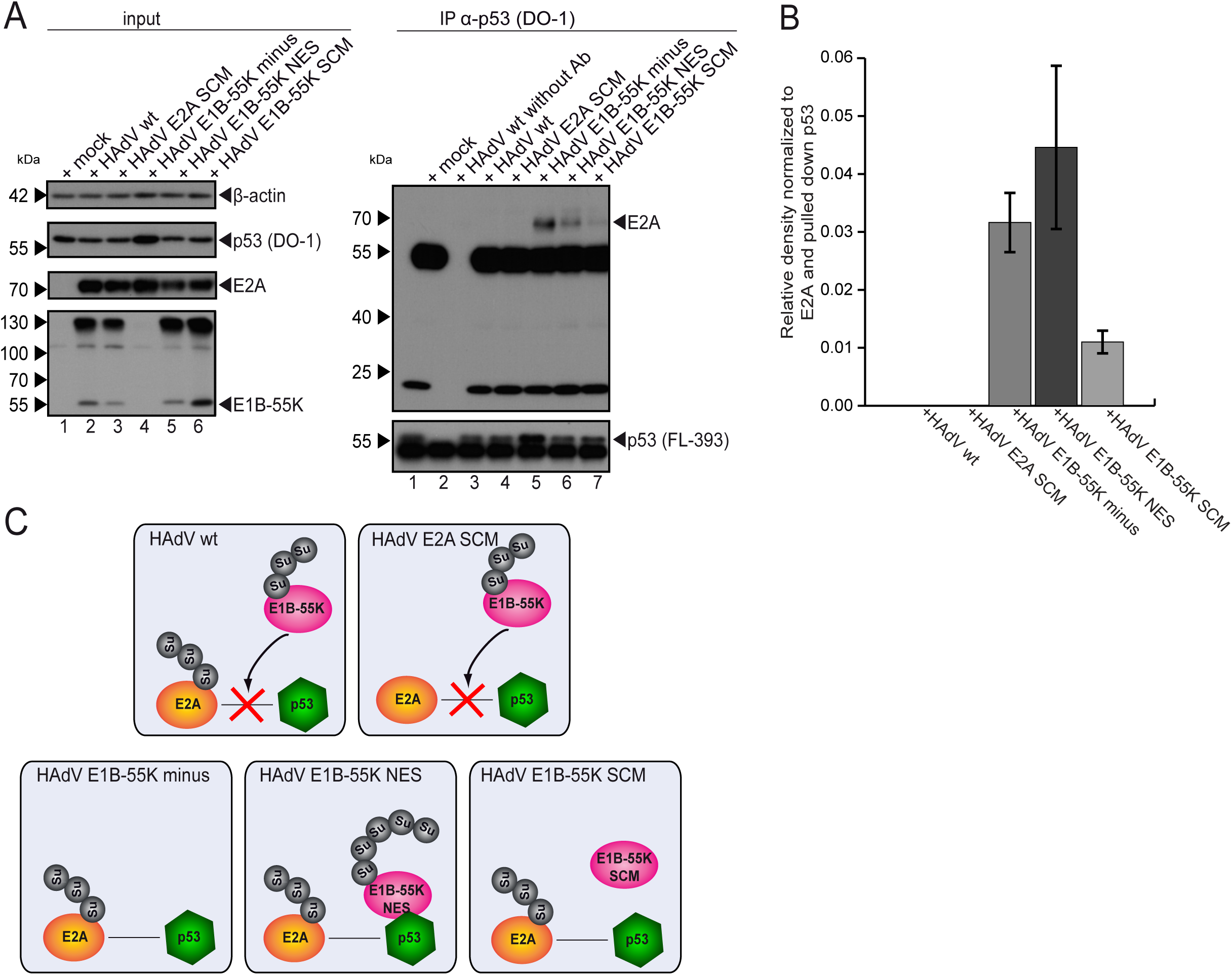
E1B-55K prevents E2A/p53 interaction during HAdV infection. **(A)** HepaRG cells were infected with HAdV wt (left panel lane 2, right panel lane 3), HAdV E2A SCM mutant (left panel lane 3, right panel lane 4), HAdV E1B-55K minus (left panel lane 5, right panel lane 6), HAdV E1B-55K NES (left panel lane 5, right panel lane 6) and HAdV E1B-55K SCM (left panel lane 6, right panel lane 7) at a multiplicity of infection of 20 FFU/cell. The cells were harvested 24 h p.i., whole-cell lysates were prepared, and immunoprecipitation analysis was performed using monoclonal mouse antibody against p53 mAb DO-1 (α-p53). Proteins were subjected to immunoblot analyses. Input levels of total-cell lysates and coprecipitated proteins were detected using mAb DO-1 (α-p53), pAb FL-393 (α-p53), mAb B6-8 (α-E2A), mAb 2A6 (α-E1B-55K) and mAb AC-15 (α-β-actin). Molecular weights in kDa are indicated on the left, relevant proteins on the right. **(B)** Densitometric analysis of E2A/p53 interaction in panel A (lanes 3, 4, 5, 6 and 7), quantified with ImageJ (version 1.45s) software, normalized to the respective levels of immunoprecipitated p53. Average values and standard deviations represented on bar plot were calculated based on three independent experiments. **(C)** Schematic representation of the data shown in panels A and B.

### E2A SUMOylation stimulates p53 degradation and SUMOylation during HAdV infection

E1B-55K cooperates with the HAdV protein E4orf6 to assemble the E3 ubiquitin ligase complex that degrades inhibitory factors for HAdV infection, including the tumor suppressor p53 (61–63). We investigated the influence of E2A SUMOylation on HAdV mediated p53 degradation by time course experiments in infected human cells. We infected the cells with HAdV wt and HAdV E2A SCM viruses. After 24 to 72 h the cells were harvested, lysed, and subjected to immunoblot analysis. HAdV E1B-55K minus and HAdV E4orf6 minus viruses were included as controls that do not degrade p53 (Fig. 4). Consistently with previous reports (61–63), p53 levels were reduced compared to mock during infection with wt virus (Fig. 4 A, lanes 2, 7 and 12). Densiometric analysis showed the reduction of p53 amount in cells infected with HAdV wt of 42.8% after 24 h p.i., 56.8% 48 h p.i. and 66.4% 72 h p.i. when compared to mock (Fig. 4 B). Infection with HAdV E2A SCM virus also led to the reduction of p53 levels compared to the mock infection (Fig 4 A, lanes 3, 8 and 13). However, densiometric analysis revealed lower reduction of p53 levels in cells infected with HAdV E2A SCM virus compared to HAdV wt infected cells, especially at 72 h p.i. The reduction of p53 levels during HAdV E2A SCM infection was 25.5% after 24 h, 39.2% after 48 h and 33.8% after 72 h when compared to mock infection (Fig. 4 B). Interestingly, we observed higher migrating bands above the p53 signals, indicating possible posttranslational modification in samples infected with HAdV wt and HAdV E2A SCM viruses, at 48 and 72 h p.i.. The signal strength of the observed bands was weaker during infection with the E2A SCM mutant virus (Fig. 4 A, lanes 2, 3, 7, 8, 12 and 13). As anticipated, degradation of p53 was not present in cells infected with HAdV E1B-55K minus and HAdV E4orf6 minus viruses (Fig. 4 A, lanes 4, 5, 9, 10, 14 and 15).

**Fig. 4.**
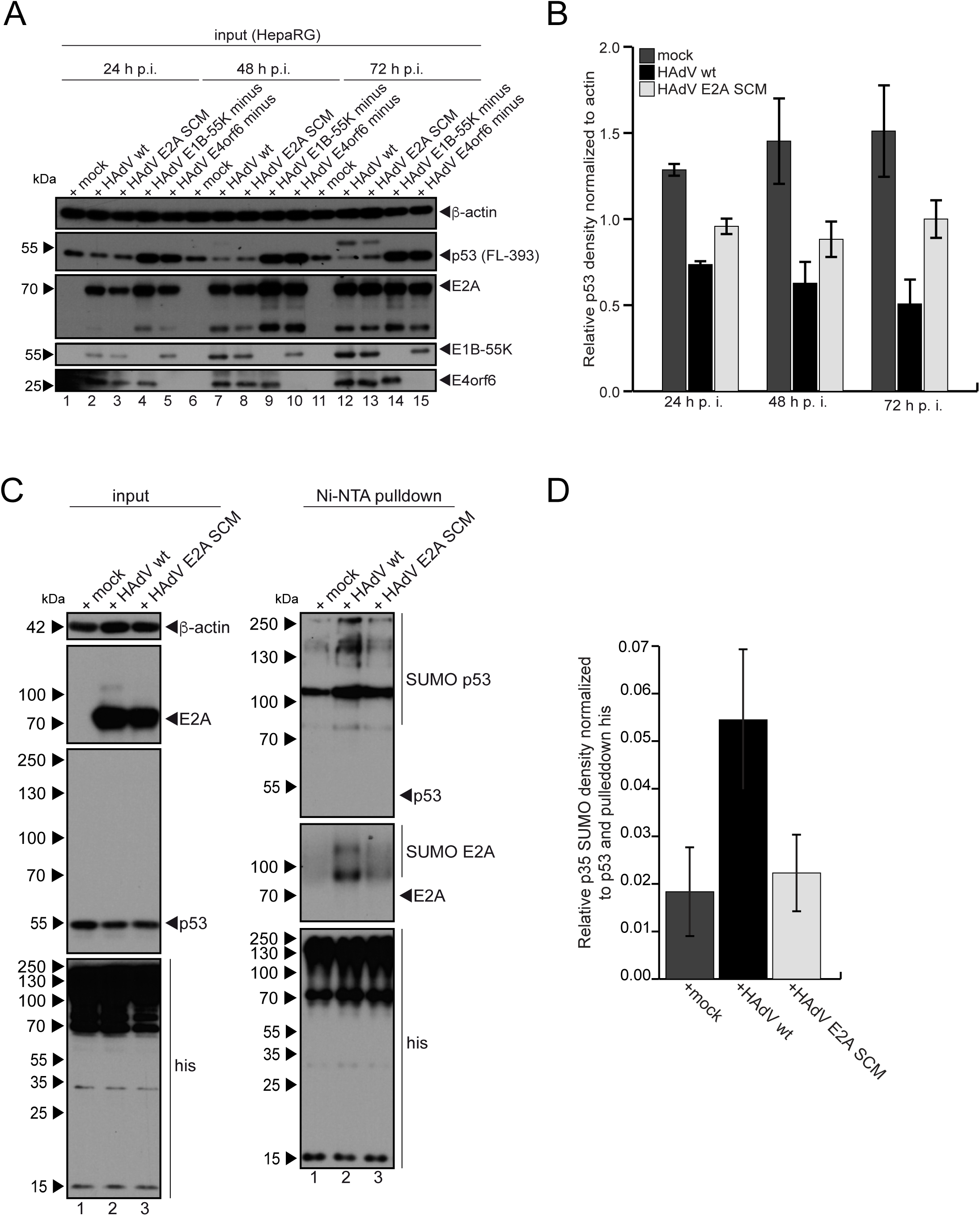
E2A SUMOylation stimulates p53 degradation and SUMOylation during HAdV infection. **(A)** HepaRG cells were infected with HAdV wt (lanes 2, 7 and 12), HAdV E2A SCM (lanes 3, 8 and 13), HAdV E1B-55K minus (lanes 4, 9 and 14) and HAdV E4orf6 minus (lanes 5, 10 and 15) viruses at a multiplicity of infection of 50 FFU/cell. The cells were harvested 24, 48 and 72 h p.i.. Total cell lysates were prepared and separated by SDS-PAGE. Immunoblotting was performed using following antibodies: pAb FL-393 (α-p53), mAb B6-8 (α-E2A), mAb 2A6 (α-E1B-55K), mAb RSA3 (α-E4orf6) and mAb AC-15 (α-β-actin). Molecular weights in kDa are indicated on the left, relevant proteins on the right. **(B)** Densitometric analysis of p53 amounts in panel A (lanes 1, 2, 3, 6, 7, 8, 11, 12 and 13), quantified with ImageJ (version 1.45s) software, normalized to the respective levels of β-actin. Average values and standard deviations represented on bar plot were calculated based on two independent experiments. **(C)** HepaRG cells stably expressing His/HA-SUMO-2 were infected with HAdV wt or HAdV E2A SCM viruses at a multiplicity of infection of 20 FFU/cell. The cells were harvested 24 h p.i., whole-cell lysates were prepared with guanidinium chloride buffer, subjected to Ni-NTA purification of 6xHis-SUMO conjugates and separated by SDS-PAGE. Input levels of total-cell lysates and Ni-NTA purified conjugates were detected with mAb B6-8 (α-E2A), pAb FL-393 (α-p53), mAb 6 His (α-6xHis-tag) and mAb AC-15 (α-β-actin). Molecular weights in kDa are indicated on the left, relevant proteins on the right. **(D)** Densitometric analysis of p53 SUMOylation levels in panel A (lanes 1, 2 and 3), quantified with ImageJ (version 1.45s) software, normalized to the respective levels of p53 and pulled down His.

E1B-55K mediated p53 SUMOylation is one of the mechanisms used by HAdV to inhibit p53 activity (58, 59). Our data reveal that E2A SUMO PTM promotes p53 degradation (Fig. 4 A, B). Additionally, we observed the higher migrating band in the p53-stained blot, indicating possible posttranslational modification of the tumor suppressor. Therefore, we investigated the influence of E2A SUMOylation on HAdV mediated p53 SUMO PTM (Fig. 4 C, D). For Ni-NTA analysis we infected HepaRG His/HA SUMO-2 cells with the HAdV wt and HAdV E2A SCM viruses at an MOI of 20 and harvested them after 24 h. We chose the MOI of 20 because after 24 h, the HAdV infection did not degrade p53 yet, and expression is still detectable. Analyses of Ni-NTA purified samples confirmed the previous finding indicating that HAdV wt infection causes the increase in p53 SUMO levels (Fig. 4 C, right panel, lane 2). p53 SUMOylation was increased by 197% in cells infected with wt virus when compared to mock infected cells (Fig. 4 D). Intriguingly, the increase in p53 SUMO levels was considerably lower in cells infected with the E2A SUMOylation deficient virus (Fig. 4 C, right panel, lane 3). Densiometric analysis, showed only 21% increase in p53 SUMOylation during infection with HAdV E2A SCM virus when compared to mock infection (Fig. 4 D). These data reveal the novel role of E2A in control of HAdV-mediated p53 degradation and SUMOylation, dependent on E2A SUMO PTM.

### E2A SUMOylation promotes E1B-55K/PML-V interaction and colocalization in HAdV infected cells

E1B-55K SUMOylates and thereby inhibits p53-dependent transactivation via its interaction with PML isoforms -IV and -V (58, 59). Our data revealed that HAdV mediated increase in p53 SUMO levels depends on E2A SUMOylation (Fig. 4 C, D). Therefore, we next investigated the influence of E2A SUMO PTM on E1B-55K/PML-V binding. As it has been previously shown that E1B-55K does not interact with PML-IV during infection (86), this PML isoform was not included in our studies. After human cells were transfected with pEYFP-PML-V and subsequently infected with wt HAdV or E2A SCM virus mutant, immunoprecipitation of GFP/YFP and staining for E1B-55K revealed previously reported interaction between the two investigated proteins during HAdV wt infection ((86), Fig. 5 A, right panel, lane 4, Fig. 5 E). However, when PML-V interaction was examined during HAdV E2A SCM infection, the amount of bound E1B-55K protein was reduced to 37.7% compared to HAdV wt infection, demonstrating an E2A SUMOylation dependent interaction between E1B-55K and PML-V (Fig. 5, right panel, lane 6, Fig. 5 B, E). During HAdV infection, E1B-55K localizes in the nucleus where it initially colocalizes with PML tracks that are induced upon HAdV infection (69, 104). As we observed the reduction of E1B-55K/PML-V binding during infection with an E2A SUMOylation deficient virus, we performed immunofluorescence studies to further substantiate this finding. The influence of E2A SUMO PTM on E1B-55K localization during infection was monitored in HepaRG cells transfected with pEYFP-PML-V and subsequently infected with HAdV wt and HAdV E2A SCM (Fig. 5 C, Fig. 5 D). To visualize that E1B-55K localized only in the nuclear matrix fraction together with PML, the cells were treated with CSK buffer before immunostaining to reduce the soluble cytoplasmic and nuclear proteins. Immunofluorescence analysis confirmed already reported data (86) showing E1B-55K colocalization with PML-V during HAdV wt infection (Fig. 5 C, panels g and h, Fig. 5 E). In contrast, the analysis of cells infected with the E2A SUMOylation deficient virus showed that E1B-55K does not colocalize with the PML-V isoform in the absence of the E2A SUMO PTM (Fig. 5 C, panels k and l, Fig. 5 E). To further substantiate the immunofluorescence findings, we determined the Pearson’s correlation coefficient for E1B-55K and PML-V that was 0.51 during HAdV wt infection confirming colocalization between the two proteins. We did not detect colocalization between the two proteins during HAdV E2A SCM infection as indicated by the Pearson’s correlation coefficient for E1B-55K and PML-V of 0.24 (Fig. 5 D).

**Fig. 5.**
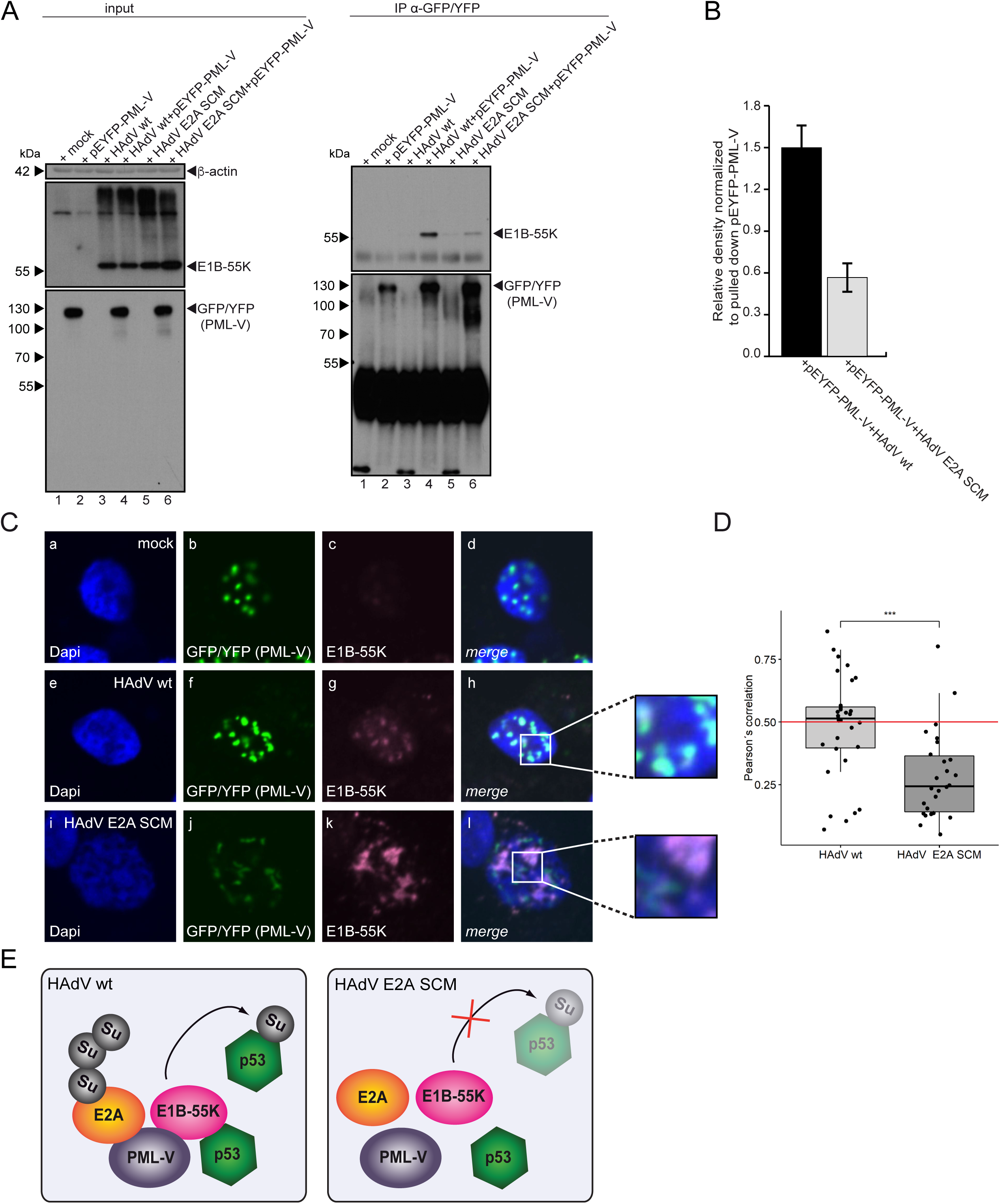
E2A SUMOylation promotes E1B-55K/PML-V interaction and colocalization. **(A)** HepaRG cells were transfected with pYFP-PML-V (lane 2) and subsequently infected with HAdV wt (lane 4) and HAdV E2A SCM mutant (lane 6) at a multiplicity of infection of 20 FFU/cell. The cells were harvested 24 h p.i., whole-cell lysates were prepared, and immunoprecipitation analysis was performed using polyclonal rabbit antibody against GFP pAb260 (α-GFP/YFP). Proteins were subjected to immunoblot analyses. Input levels of total-cell lysates and coprecipitated proteins were detected using mAb 2A6 (α-E1B-55K), pAb260 (α-GFP/YFP) and mAb AC-15 (α-β-actin). Molecular weights in kDa are indicated on the left, relevant proteins on the right. **(B)** Densitometric analysis of E1B-55K interaction with PML-V in panel A (lanes 4 and 6), quantified with ImageJ (version 1.45s) software, normalized to the respective levels of immunoprecipitated YFP-PML-V. Bar plot represents average values and standard deviations calculated based on data obtained in three independent experiments. **(C)** HepaRG cells were transfected with 3 µg of pYFP-PML-V and subsequently infected with HAdV wt or HAdV E2A SCM mutant at a multiplicity of infection of 20 FFU/cell, 4 h post transfection. Prior to fixation with 4% PFA, 24 h p.i., the cells were treated with CSK buffer to remove soluble cytoplasmic proteins and loosely held nuclear proteins. Fixed cells were double labeled with pAb260 (α-GFP/YFP) and mAb 2A6 (α-E1B-55K). Primary Abs were detected with Alexa488 (GFP/YFP, green) and Alexa647 (E1B-55K, pink) conjugated secondary Abs. Nuclei were stained by Dapi. Anti-GFP/YFP (green; panels b, f, j) and E1B-55K (pink; panels c, g, k), staining patterns were analyzed in 30 cells. Overlays of the single images (merge) are shown in panels d, h, l. **(D)** Pearson’s correlation coefficient for YFP-PML-V stained by pAb260 (α-GFP/YFP) antibody and E1B-55K stained by mAb 2A6 (α-E1B-55K) antibody in cells infected with HAdV wt or HAdV E2A SCM was determined using the Volocity software. For each cell, Pearson’s correlation was calculated for colocalization of YFP-PML-V with E1B-55K. The distributions of the respective correlation coefficients are displayed in a box plot. Values above 0.5 are considered colocalization. Statistical significances for differences in determined YFP-PML-V/E1B-55K Pearson’s correlation coefficients between HAdV wt and HAdV E2A SCM infected cells, were determined using two-sided Mann-Whitney-U test. ***, P < 0.001. **(E)** Schematic representation of the data shown in panels A, B, C and D.

### HAdV inhibits p53/DNA binding on an E2A SUMOylation dependent manner

According to our data, E2A SUMOylation is required for efficient degradation of p53 (Fig. 4 A, B). Furthermore, we showed that SUMOylated E2A promotes HAdV-mediated p53 SUMOylation (Fig. 4 C, D) by regulating E1B-55K binding and colocalization with PML-V (Fig. 5). E1B-55K acts as a transcriptional repressor of p53-dependent genes (105). E1B-55K interaction and colocalization with PML-NBs is tightly linked to the inhibition of p53 activity and thereby oncogenic properties of the viral factor (58, 59). As p53 achieves its activity through binding to promotor regions of its target genes (5, 6), we investigated the influence of E2A SUMOylation on p53 binding to DNA during HAdV infection. ChIP-seq reads of mock infected cells or cells infected with HAdV wt and HAdV E2A SCM viruses were aligned to the human genome (hg38), and p53 binding sites (p53 peaks) were identified using MACS2 (89) (Fig. 6 A, B). Peaks of all three conditions were merged into a combined p53 peak set using bedtools (90) and annotated with the annotatePeaks.pl script of the HOMER sequencing analysis package (Fig. 6 B). We found numerous p53 binding sights on the host genome (Fig. 6 A, Suppl. Table 1). Our analysis revealed overall reduction of p53 binding to DNA in cells infected with HAdV wt virus compared to mock infected cells, that was not detected in HAdV E2A SCM infected cells (Fig. 6 A, B). We found 95 differential annotated peaks in mock infected cells compared to HAdV wt infection, 99 differential annotated peaks in HAdV E2A SCM infected cells compared to HAdV wt infection and only 13 differentiated annotated peaks between HAdV E2A SCM and mock (Fig. 6 C). Furthermore, we detected reduction of p53 binding within the regulatory regions of key p53-targeted genes, such as *MDM2* and *CDKN1A* in cells infected with HAdV wt that was absent during HAdV E2A SCM infection (Fig. 6 D). Data shown here indicate the essential role of E2A SUMO PTM in HAdV mediated inhibition of p53 binding to DNA.

**Fig. 6.**
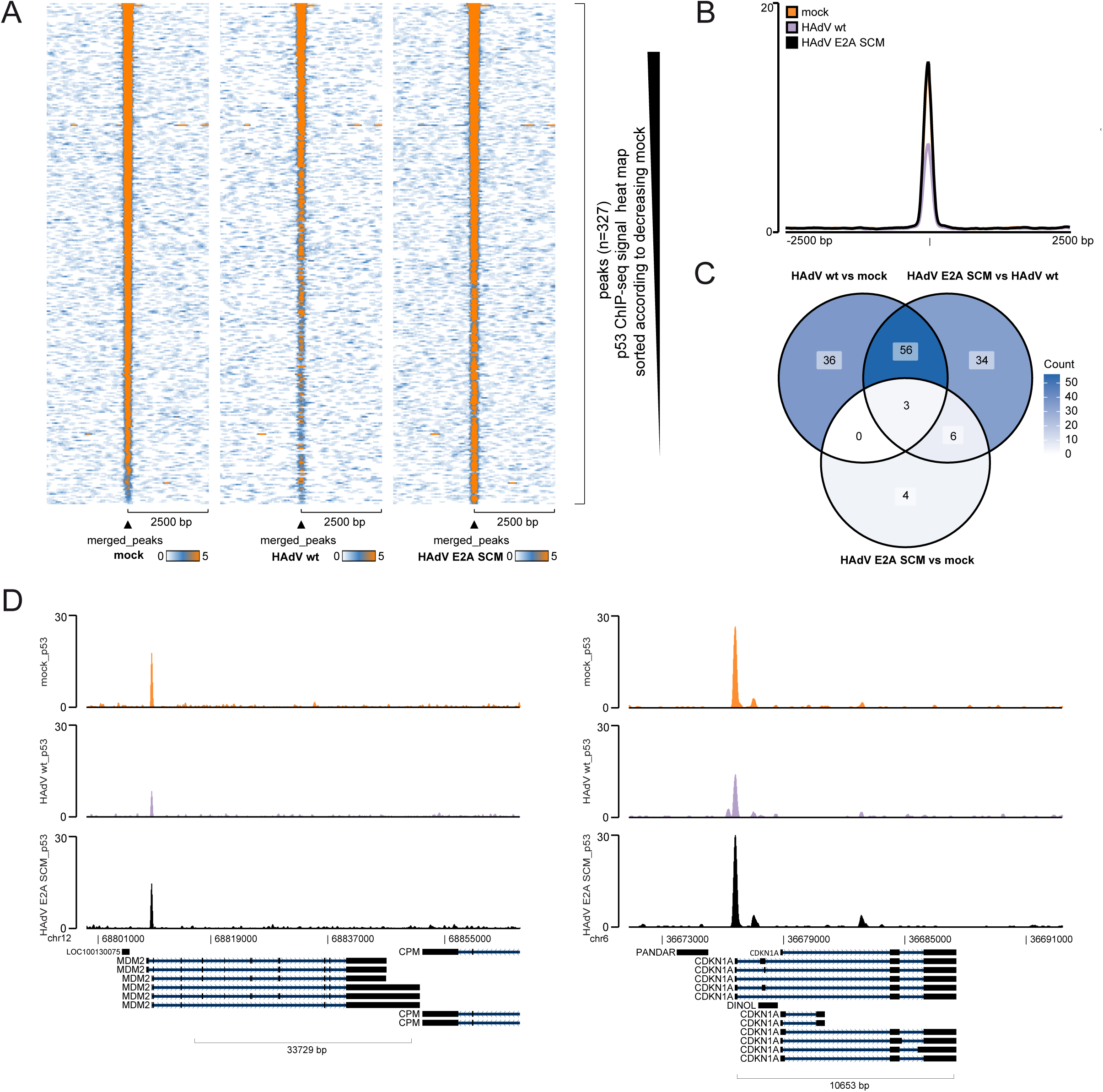
HAdV inhibits p53/DNA binding on an E2A SUMOylation dependent manner. HepaRG cells were mock infected or infected either with HAdV wt or HAdV E2A SCM virus at a multiplicity of infection of 20 FFU/cell and harvested for ChIP analysis 24 h p.i. ChIP-seq binding profiles of p53 were obtained via alignment to the human genome (hg38). **(A)** Illustration of p53 ChIP-seq signal with heat map normalized to RPGC (reads per genomic content). The normalized p53 ChIP signal intensity for mock, HAdV wt or HAdV E2A SCM infected cells is displayed in a color gradient from light blue (0, weak) to orange (5, strong). **(B)** The profile plot displays the mean signal of p53 within its respective peak set. **(C)** Venn diagram representing ChIP-seq peaks differentially regulated in mock, HAdV wt and HAdV E2A SCM infected cells associated to annotated genes. **(D)** Visualization of detected peaks in two representative regulatory elements of p53 target gene regions (MDM2 and CDKN1A) in mock, HAdV wt and HAdV E2A SCM infected cells. The y-axes indicate the respective ChIP-seq signal, normalized to reads per million per 1 kb.

### E2A SUMOylation inhibits p53-mediated transcription during HAdV infection

Our data showed an E2A SUMO PTM role in degradation and SUMOylation of p53 (Fig. 4). Furthermore, we showed that SUMOylated E2A regulates E1B-55K interaction and colocalization with PML-V (Fig. 5), required for the inhibition of p53 activity (58, 59). Based on these findings, our data also showed that E2A SUMOylation controls HAdV mediated inhibition of p53/DNA binding (Fig. 6), necessary for activation of its target genes (5, 6). Therefore, we investigated the influence of E2A SUMOylation on HAdV-mediated modulation of p53-dependent transcriptional activation. We evaluated the effect of E2A SCM mutations on HAdV infection induced host transcriptome changes with RNAseq approaches. Sequencing libraries were prepared in triplicates for cells infected with HAdV wt and HAdV E2A SCM viruses and analyzed comparing the respective infected cell lines and mock infected cells. Our data revealed 9497 differentially expressed transcripts of 6078 genes in HAdV wt infected cells compared to mock, while 1687 transcripts of 1276 genes were differentially expressed, comparing HAdV wt and HAdV E2A SCM infected cells (Fig. 7 A, Suppl. Table 2). Functional GO term analysis of biological processes in HAdV wt infected cells compared to mock revealed differential expression of transcripts mostly involved in protein synthesis (eg. *cytoplasmic translation, ribosome biogenesis, rRNA metabolic process, RNA splicing*) and cell cycle regulation (eg. *regulation of chromosome organization, regulation of cell cycle phase transition, mitotic cell cycle phase transition*) (Fig. 7 B, left panel). Transcripts differentially expressed in HAdV E2A SCM infected cells compared to HAdV wt infection (Fig. 7 B, left panel) were, on the other hand, mostly involved in cytoskeleton organization (eg. *regulation of supramolecular fiber organization, actin filament organization, positive regulation of cytoskeleton organization*), nucleotide metabolism (eg. *ribose phosphate metabolic process, ribonucleotide metabolic process, nucleoside triphosphate metabolic process*) and protein homeostasis (eg. *regulation of protein catabolic process, protein polymerization, regulation of protein−containing complex assembly*). The term ‘‘*intrinsic apoptotic signaling pathway*’’ that was also enriched in our analysis indicated deregulation of p53 activity as this protein is considered master regulator of apoptotic process (2–4). This and the findings so far shown in this study indicate an E2A SUMOylation mediated regulation of p53 activity. Thus, this directed further investigation with a focus on transcripts of p53 target genes, as reported by Fischer and colleagues (2). Among the statistically significantly differentially expressed transcripts shown in volcano plot, we highlighted transcripts of p53 target genes that were previously described (2) (Fig. 7 C). Our data revealed 337 p53 target gene transcripts of 198 genes differentially expressed between noninfected and HAdV wt infected cells (Fig. 7 D. Suppl. Table 3). Intriguingly, 54 transcripts of 43 genes were differentially expressed, comparing HAdV wt and HAdV E2A SCM infected cells (Fig. 7 D). We further analyzed these transcripts. The heat map represents the top 50 transcripts of p53 target genes differentially expressed in HAdV wt and HAdV E2A SCM infection. Most transcripts were upregulated in HAdV E2A SCM infection compared to HAdV wt infection. However, transcript expression of some genes (*CCDC51*, *FOSL1*, *ITGA3*, *TP53*, *MR1*, *VCAN*), was downregulated when comparing HAdV E2A SCM infected cells with HAdV wt infected cells, while for *LRPAP1* transcripts were heterogeneously expressed showing both, up-and down regulated expression (Fig. 7 E). Bar plot represents log2 fold changes (log2FC) for expression of top 50 transcripts of p53 target genes differentially expressed when comparing HAdV E2A SCM to HAdV wt infected cells and HAdV wt infected cells to non-infected cells (Fig. 7 F). In concordance with previous reports, most of p53 target gene transcripts were downregulated in HAdV wt infected cells compared to non-infected cells (106, 107). However, vast majority of p53 target gene transcripts differentially expressed between HAdV E2A SCM and HAdV wt infected cells (eg. *BBC3*, *CDKN1A*, *GADD45A*) were upregulated in the absence of efficient E2A SUMOylation. Our data also showed several downregulated transcripts in HAdV E2A SCM compared to HAdV wt infected cells, whose expression was upregulated in HAdV wt infected cells when compared to mock infection (*CCDC51*, *FOSL1*, *ITGA3*, *TP53*, *LRPAP1.2*). Additionally, *MR1* and *VCAN* transcription was reduced in HAdV wt infected cells compared to non-infected cells but even more reduced in HAdV E2A SCM infected cells compared to HAdV wt infected cells (Fig. 7 F). In sum, our findings clearly demonstrate the requirement for E2A SUMOylation for efficient inhibition of p53 activity during HAdV infection.

**Fig. 7.**
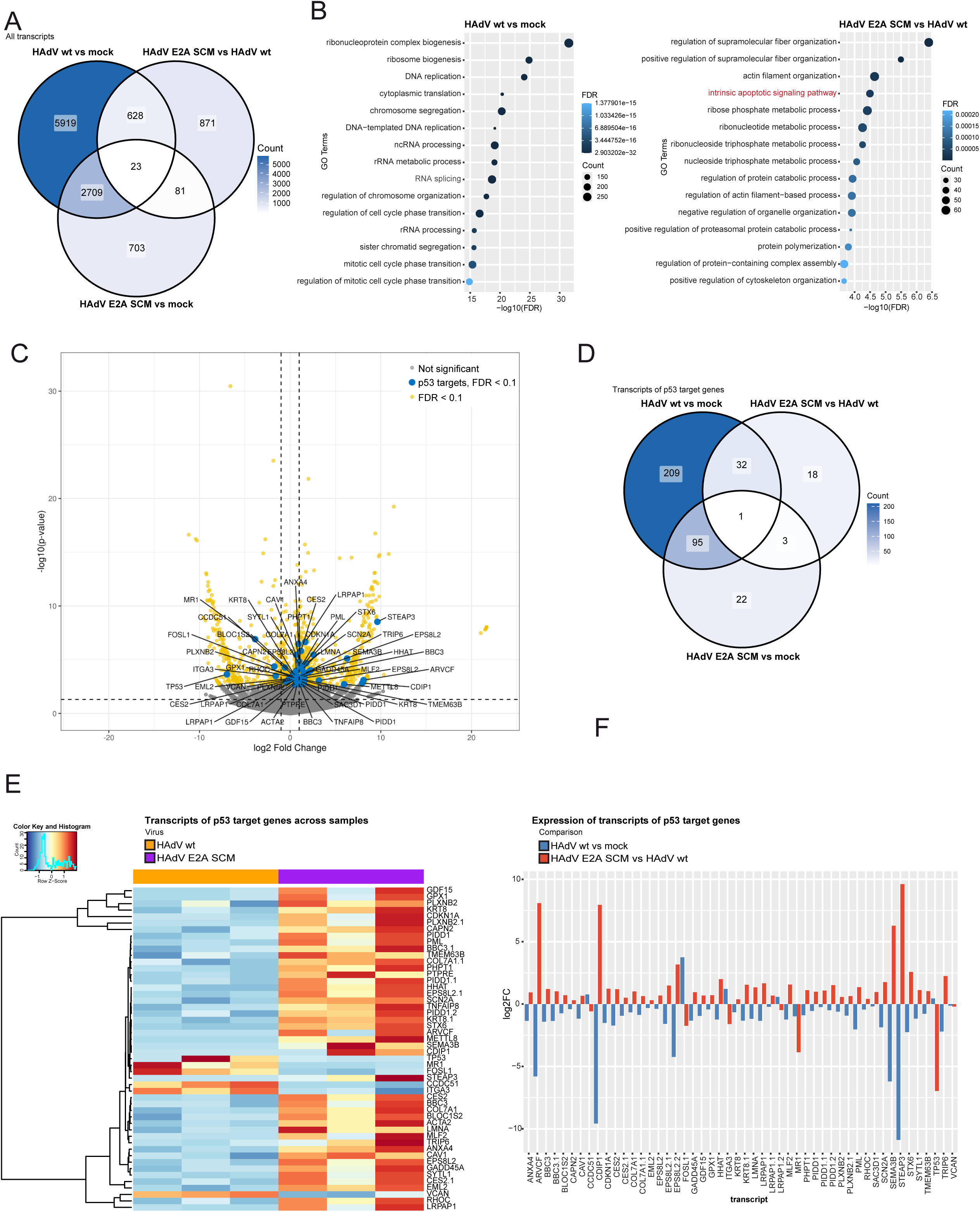
HAdV inhibits transcript expression of specific p53 target genes using E2A SUMOylation. HepaRG cells were mock infected or infected either with HAdV wt or HAdV E2A SCM virus at a multiplicity of infection of 20 FFU/cell and harvested 24 h p.i. Total RNA was extracted and subjected to Illumina RNA sequencing. Differential transcript expression was computed and p-values were corrected for multiple comparisons using a cutoff of FDR < 0.1. **(A)** Venn diagram showing differentially expressed transcripts detected with Illumina sequencing in mock, HAdV wt and HAdV E2A SCM infected cells. **(B)** Gene ontology (GO) term analysis of all differentially regulated transcripts between HAdV wt and mock (left), and HAdV E2A SCM and HAdV wt (right) infected cells. Dot plots show the top 15 GO terms for biological processes ranked by adjusted p-value (FDR < 0.05) following analysis of all differentially expressed transcripts. The cluster term “*intrinsic apoptotic signaling pathway*” is highlighted in red. **(C)** Volcano plot highlighting the distribution of transcripts differentially expressed in HAdV E2A SCM compared to HAdV wt infected cells. Significantly differentially expressed transcripts (FDR < 0.1) are colored yellow. Significantly differentially expressed transcripts of prominent p53 target genes reported by Fischer et al. (2) are colored blue. Not significant transcripts are colored gray. The –log10 p-values for each transcript are plotted on the *y* axis. The vertical dashed line indicates 2 fold increase or decrease of transcript expression in HAdV E2A SCM compared to HAdV wt infected cells. Horizontal dashed line signifies p-value of 0.05. The complete list of underlying transcripts is provided in (Suppl. Table 2). **(D)** Venn diagram showing differentially expressed transcripts of prominent p53 target genes reported by Fischer et al. (2) detected with Illumina sequencing in mock, HAdV wt and HAdV E2A SCM infected cells. **(E)** Heatmap of the top 50 of p53 target gene transcripts, differentially regulated in HAdV E2A SCM compared to HAdV wt infected cells, (FDR < 0.1 and log2 fold change > 0). Shown are row Z-scores and counts of normalized reads of indicated transcripts. **(F)** Bar plot represents log2fold changes (log2FC) of the top 50 significantly differentially expressed transcripts of p53 target genes between HAdV wt and HAdV E2A SCM infected cells. Log fold change was computed for each transcript comparing HAdV E2A SCM infected cells to HAdV wt infected cells (HAdV E2A SCM vs HAdV wt) and HAdV wt infected cells to not-infected cells (HAdV vs mock).

### HAdV-mediated apoptosis inhibition during infection is E2A SUMOylation dependent

Our data showed that E2A SUMOylation controls HAdV mediated inhibition of p53 activity during infection (Fig. 7). As one of major p53 functions is control of host cells apoptotic process (108), we further investigated the role of E2A SUMOylation on HAdV-mediated inhibition of apoptosis.

We performed KEGG pathway analysis of statistically significantly expressed transcripts in HAdV E2A SCM infected cells compared to HAdV wt infected cells and detected *‘‘Apoptosis’’* as most significantly enriched pathway (Fig. 8 A). We further focused on transcripts of genes belonging to this pathway, according to KEGG (109–111) (Suppl. Table 4). We detected 111 transcripts of 64 genes involved in control of apoptotic process differentially expressed between not-infected and HAdV wt infected cells and 45 transcripts of 30 genes differentially expressed between HAdV E2A SCM and HAdV wt infected cells. Our generated heat map shows differentially expressed transcripts of genes involved in control of apoptosis in cells infected with HAdV E2A SCM compared to HAdV wt infected cells (Fig. 8 B). As transcripts of p53 target genes (Fig. 7, Suppl. Table 3), most apoptosis related transcripts were upregulated in HAdV E2A SCM infection compared to HAdV wt infection (Suppl. Table 4). Only a minor portion of differentially expressed transcripts was downregulated in HAdV E2A SCM infected cells compared to HAdV wt infected cells (*CTSF*, *TNFRSF1A*, *TP53*, *TUBA4A*). We also detected heterogenous expression of certain transcripts that were both up and down regulated (*ACTB*, *BAD*, *MAP2K2*, *PTPN13*) (Fig. 8 B). Bar plot represents log2 fold changes (log2FC) for expression of apoptosis related transcripts differentially expressed in HAdV E2A SCM compared to HAdV wt infected cells and in HAdV wt infected cells compared to non-infected cells (Fig. 8 C). Majority of transcripts was downregulated in HAdV wt infected cells compared to non-infected cells, as previously reported (106). On the other hand, apoptosis related transcripts were mostly upregulated in HAdV E2A SCM infected cells compared to HAdV wt infected cells (Fig. 8 C, Suppl. Table 4). Our data also showed several downregulated transcripts comparing HAdV E2A SCM to HAdV wt infected cells, whose expression was upregulated in HAdV wt infected cells when compared to mock infection (*CTSF*, *TNFRSF1A*, *TP53*, *TUBA4A*, *ACTB*, *BAD*, *MAP2K2*, *PTPN13.1*) (Fig. 8, Suppl. Table 4). This data clearly indicate the role of E2A SUMOylation in HAdV mediated manipulation of apoptosis. Therefore, we determined host cell apoptosis activation by measuring caspase-3/7 activity in non-infected mock cells or cells infected with HAdV wt or HAdV E2A SCM viruses. We infected HepaRG (MOI 50) and A549 (MOI 20) cells and treated them 48 h p.i. with actinomycin D for 4.5 h to induce apoptosis (Fig. 8 D, E). Luminescence measurements in both examined cell lines confirmed previous reports (112) showing that HAdV wt infection inhibits apoptosis (Fig. 8 D, E). Caspase-3/7 activity was decreased by 44% in HepaRG and 185% in A549 cells infected with wt virus when compared to mock infected cells (Fig. 8 D, E). Intriguingly, the decrease in caspase-3/7 activity was significantly lower in cells infected with the E2A SUMOylation deficient virus as luminescence measurements showed decrease of 28% in HepaRG and 138% in A549 cells during infection with HAdV E2A SCM virus when compared to mock infection (Fig. 8 D, E). To gain better insight into the role of E2A SUMOylation in inhibition of apoptosis during HAdV infection, we mapped the most significant differentially expressed transcripts of each gene on apoptosis KEGG pathway chart using ggKEGG (113). We mapped the comparisons between HAdV E2A SCM and HAdV wt (Fig. 8 F), and HAdV wt and mock infected cells (Suppl. Fig. 2). Significantly upregulated transcripts are labeled in red, downregulated in blue.

**Fig. 8.**
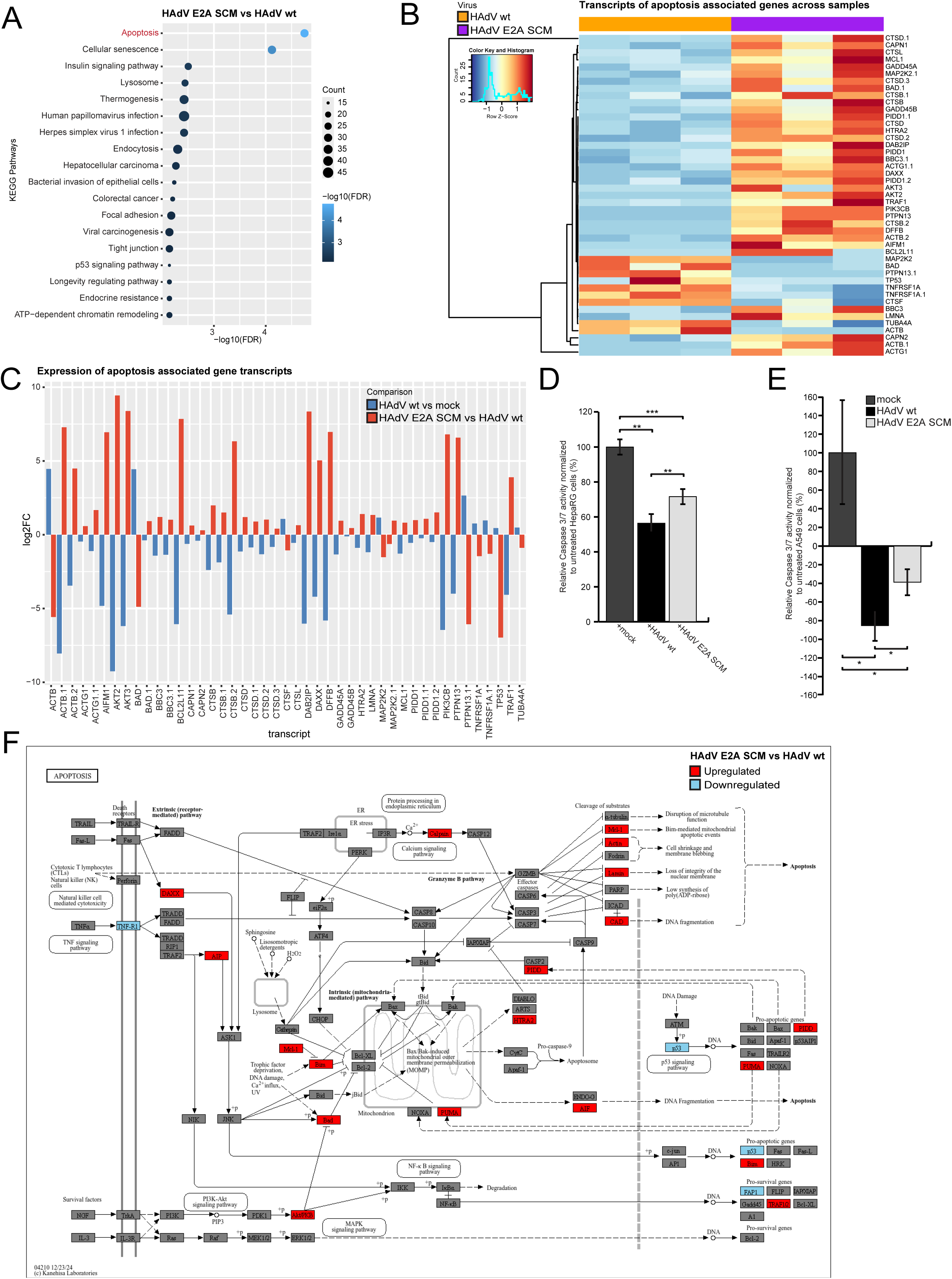
HAdV inhibits apoptosis on an E2A SUMOylation dependent manner. HepaRG cells were mock infected or infected either with HAdV wt or HAdV E2A SCM virus at a multiplicity of infection of 20 FFU/cell and harvested 24 h p.i. Total RNA was extracted and subjected to Illumina RNA sequencing. Differential transcript expression was computed and p values were corrected for multiple comparisons using a cutoff of FDR < 0.1. **(A)** KEGG pathway analysis of all differentially regulated transcripts between HAdV E2A SCM and HAdV wt infected cells. Dot plot shows the enriched KEGG pathways ranked by adjusted p-value (FDR < 0.05) following analysis of all differentially expressed transcripts. The cluster pathway “*Apoptosis*” is highlighted in red. **(B)** Heatmap of the gene transcripts associated with apoptosis pathway, differentially regulated in HAdV E2A SCM compared to HAdV wt infected cells, (FDR < 0.1 and log_2_ fold change > 0). Shown are the top 50 significantly differentially expressed transcripts with corresponding row Z-scores and counts of normalized reads. **(C)** Bar plot represents log2fold changes (log2FC) of the top 50 significantly differentially expressed transcripts of apoptosis pathway associated genes, between HAdV E2A SCM and HAdV wt infected cells. Log fold change was computed for each transcript comparing HAdV E2A SCM infected cells to HAdV wt infected cells (HAdV E2A SCM vs HAdV wt) and HAdV wt infected cells to not-infected cells (HAdV vs mock). **(D)** HepaRG and A549 **(E)** cells were HAdV wt, HAdV E2A SCM or mock infected at a multiplicity of infection of 50 FFU/cell **(D)** or at a multiplicity of infection of 20 FFU/cell **(E)**. 48 hp.i. HepaRG and A549 cells were treated with 0.5 µM **(D)** or 10µM **(E)** Actinomycin D for 4.5 h to induce apoptosis. Caspase-Glo 3/7 Reagent (Promega) was added directly to the cells in 96-well plates and incubated for 1 h before measuring luminescence. Recorded luminescence values for treated cells were normalized by subtracting the luminescence values measured in non-infected cells equally treated with Actinomycin D solvent (DMSO). Average values and standard deviations represented on bar plots were calculated based on three independent experiments. Statistical significance was determined using Welch’s t-test (*, P < 0.05, **, P < 0.01). Average luminescence values were shown as percentage of mock infected cells. **(F)** Data obtained by transcriptomic analysis of differentially expressed transcripts in HAdV E2A SCM infected cells compared to HAdV wt infected cells was rendered on KEGG apoptosis pathway graph using ggKEGG (113). Most significantly expressed transcript of each gene (FDR < 0.1) was rendered on the KEGG apoptosis pathway graph. Red color signifies upregulation, blue downregulation.

**Fig. 9.**
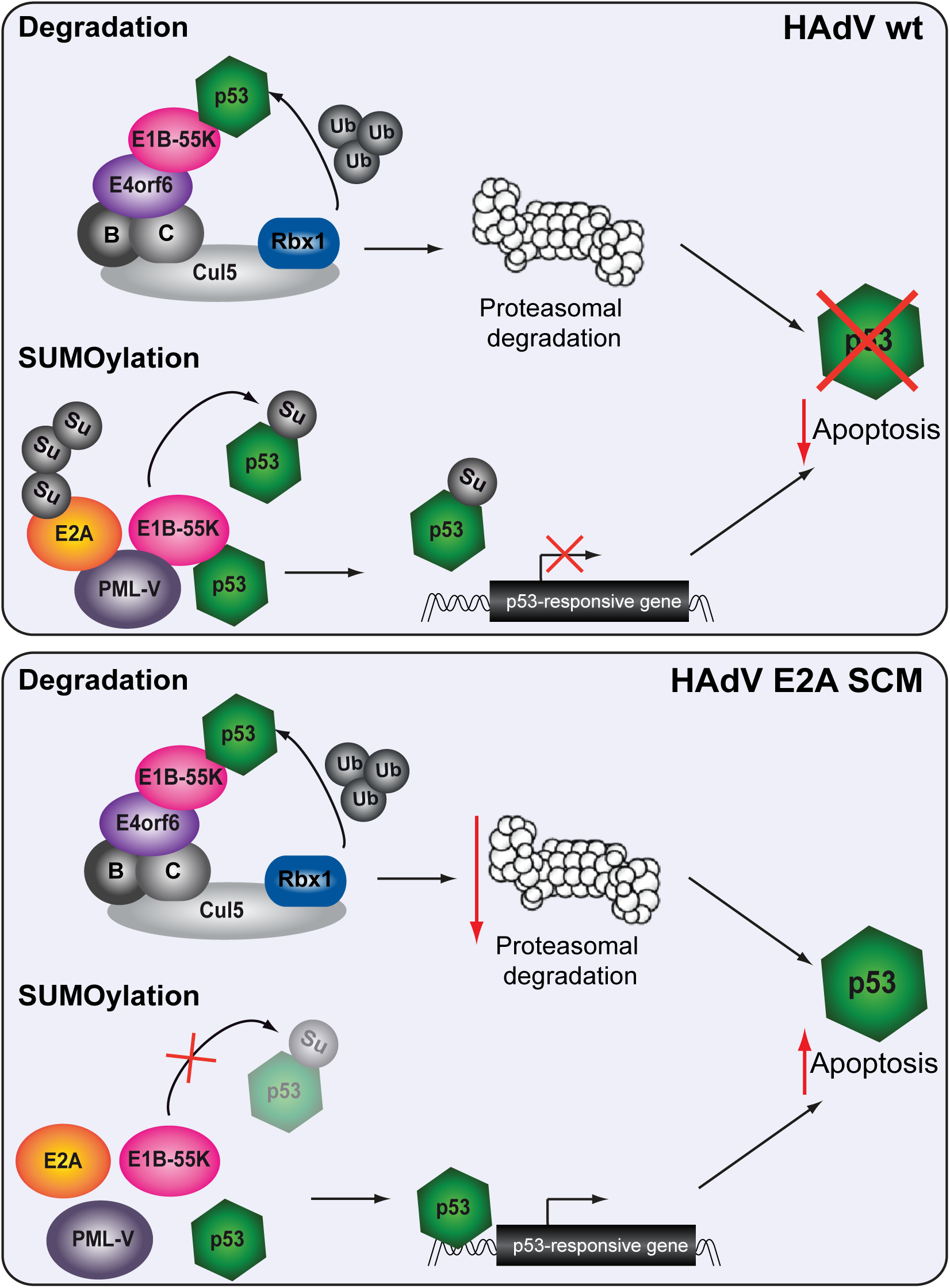
Schematic representation of the E2A SUMOylation mediated inhibition of p53 activity. Proposed model links E2A SUMOylation with the inhibition of the p53 activity during HAdV infection. The model shows the role of E2A SUMO PTM in proteasomal degradation and inhibition of p53. E2A SUMOylation promotes E2A/PML binding and E1B-55K interaction and colocalization with PML-V thereby causing the increase in p53 SUMOylation and subsequent inhibition of the p53 binding to p53 responsive elements. This suppresses p53 mediated transcriptional activation, leading to the inhibition of apoptosis.

## DISCUSSION

The activation of the tumor suppressor p53 leads to growth inhibition by induction of cell cycle arrest or apoptosis (2–4). Besides its key role in tumor suppression, p53 is also involved in antiviral innate immunity by both inducing apoptosis and enforcing the type I interferon response (11). Therefore, numerous viruses have developed intricate mechanisms to counteract its activity. In particular, p53 function is repressed during Human Cytomegalovirus (HCMV) infection (12, 13), and Human immunodeficiency virus-1 (HIV-1) and Human papillomavirus (HPV) infections trigger p53 degradation (15, 20). Interestingly, during Herpes simplex virus type 1 (HSV-1) infection p53 acts both as positive and negative factor (17), while Influenza A infection benefits from p53 functions (114). HAdV and SV-40 infections lead cells to the S phase, resulting in p53 accumulation, which causes apoptosis if p53 activity is not disrupted (115–117). Therefore, both viruses employ different strategies to inhibit p53 and assure efficient viral replication (14, 16, 18, 19).

One of the HAdV strategies to inactivate p53 by early protein E1B-55K is to affect p53 subcellular localization to regulate the tumor suppressor activity (99–101). In detail, E1B-55K sequesters p53 into perinuclear bodies, however during HAdV infection p53 was also detected in E1B-55K containing sites that resemble to HAdV RCs (56). Nevertheless, a clear association of p53 with HAdV RCs has not been reported up to now. Based on our findings, we conclude that p53 localizes to HAdV RCs during infection on an E1B-55K- and E2A SUMOylation-independent manner. As E2A represents the marker protein for HAdV RCs (68), we hypothesize that E2A might recruit p53 to RCs. Nevertheless, we could not detect the E2A/p53 interaction during HAdV wt or HAdV E2A SCM infection. We conclude that functional E1B-55K blocks this binding, to presumably protect HAdV RCs from negative effects of p53. In the absence of E1B-55K, that directly binds the tumor suppressor (51, 52), there is no barrier for the interaction between E2A and p53. When E1B-55K NES is inactive, E1B-55K is highly SUMOylated and accumulates at PML-NBs and the periphery of viral RCs (54). This offers a plausible explanation for E2A/p53 binding during infection with HAdV E1B-55K NES virus, as the portion of p53 that would be in cytoplasmic inclusions during wt infection can target HAdV RCs. E1B-55K SCM mutation abrogates E1B-55K nuclear localization (84), and therefore the barrier between p53 localizing in RCs and with E2A.

Besides the change in localization, HAdV inhibits p53 activity by proteasomal degradation via the E3 ubiquitin ligase complex assembled by E1B-55K and E4orf6 (61–63). Based on our data showing that the E2A SCM mutant retarded p53 degradation during infection, we conclude that E2A SUMO PTM promotes p53 inhibition mediated by the HAdV E3 ubiquitin ligase complex. HAdV also inhibits the p53 transcriptional activity by E1B-55K mediated p53 SUMOylation, dependent on the interaction of the viral factor with PML-IV and PML-V (58, 59, 105). Our findings reveal that E2A SUMO PTM is required for the induction of p53 SUMOylation during infection. We conclude that E2A SUMOylation underlays E1B-55K interaction and colocalization with PML-V. Therefore, the observed absence of p53 SUMOylation during infection with E2A SCM mutant virus is a consequence of the reduction of E1B-55K/PML-V binding in cells infected with the E2A SUMOylation deficient virus. Furthermore, we hypothesize that recently reported E2A SUMOylation dependent E2A/PML interaction and PML tracks localization next to HAdV RCs (77) could be the prerequisite for the E1B-55K/PML-V binding and colocalization. As the transforming activity of E1B-55K highly depends on its PML-NBs localization (59) we hypothesize that the infection with E2A SUMOylation deficient virus would exhibit impaired transforming potential in non-permissive cells.

Previous studies showed that p53 performs its functions through binding to promotor regions of its target genes (5, 6). Our data showed downregulation of p53 binding to DNA caused by HAdV wt infection, absent in cells infected with E2A SUMOylation deficient virus. This goes in line with our transcriptome analysis of infected cells that revealed a subset of p53 dependent transcripts downregulated by the wt virus but not by the E2A SUMOylation deficient virus. Based on this, we conclude that SUMOylated E2A supports HAdV-mediated inhibition of p53 transactivation. Our data also revealed a small group of transcripts that were upregulated by the wt virus but not by the E2A SCM mutant virus. This can be explained by dual regulation in response to p53 as in case of FOSL1 (118). In accordance with previous reports, TP53 expression was upregulated in wt virus infected cells (106, 107). E1A binds pRb (retinoblastoma protein) related proteins enabling their dissociation from the transcription factor E2F family, leading to the increased expression of E2F responsive genes, including TP53, required for the S-phase entry (19, 41–44). E1A localizes in PML-NBs during infection (75). PML tracks are not located next to HAdV RCs when E2A is not SUMOylated (77). Based on this we hypothesize that downregulation of TP53 expression in HAdV E2A SCM infected cells, that our data revealed, could arise from E1A inability to efficiently perform its functions. This assumption goes in line with findings reported here, demonstrating the need for E2A SUMOylation for E1B-55K optimal localization and function. However, further studies are needed to confirm this hypothesis.

Even though our data clearly indicate E2A SUMOylation mediated control of p53 activity, introduction of SCM mutations in E2A does not affect the expression of all p53 dependent transcripts observed during HAdV wt infection. Therefore, we assume that the virus uses the E2A SUMO PTM to inhibit only specific p53 functions. Based on our data, we conclude that E2A SUMOylation controls HAdV mediated inhibition of apoptosis. In accordance with previously reported findings, most of the apoptosis related transcripts were downregulated in wt infected cells (106). However, we found significant group of apoptosis related transcripts that were upregulated in the absence of E2A SUMOylation, compared to wt infection. This was further supported by measurement of apoptotic activity that was higher in cells infected with E2A SUMOylation deficient virus. The inability to fully inhibit p53 activity and apoptotic response from the host cell offers an additional explanation for the impaired HAdV progeny production during infection with the E2A SUMOylation deficient virus, recently reported by our group (77).

HAdV employs additional mechanism mediated by E1B-55K to inhibit p53 activity leading to cell transformation. E1B-55K interacts with Sp100A prior to SUMOylation and sequesters the host factor into the insoluble matrix of the nucleus or into cytoplasmic inclusions. E1B-55K/Sp100A complex recruits p53 and inhibits its transcription activating properties (60). Both, interaction of E1B-55K with PML and Sp100A, are dependent on E1B-55K SUMO PTM (60, 86). Furthermore, the interaction of E2A with these two cellular factors is also E2A SUMOylation dependent (77). Based on these reports, it is tempting to speculate that E2A employs the mechanism independent on E1B-55K, to inhibit p53 through Sp100A and PML interaction. On the other hand, E2A SUMOylation could also be the prerequisite for E1B-55K interaction with Sp100A leading to the p53 inhibition.

HAdVs are thoroughly researched and used in the context of their oncolytic potential as well as vaccine and gene therapy vectors (30–33, 119). Here we demonstrate a so far unknown role of E2A SUMO PTM in HAdV-mediated inhibition of p53 activity and apoptosis, indicating the possible oncogenic potential of E2A. Introduction of E2A SCM mutations into adenoviral vectors that are used as therapeutics should thus be explored as a viable option to improve their safety and efficacy. Oncolytic AdV therapy approaches use the ability of the virus to cause the cell lysis and engineered tumor selectivity is built on the knowledge on HAdV-mediated inhibition of DNA damage response and apoptosis. Tumor selectivity can be accomplished by deleting parts of viral genes (120). It was reported that E1B-55K deleted virus, dl1520/ONYX-015, induces high p53 levels, expected to limit viral replication in healthy cells but not in p53 mutant tumor cells (121). Nevertheless, high p53 levels in dl1520/ONYX-015 infected human cells do not activate all p53 transcriptional targets (50). As this limits the treatment success, it was proposed that the deletion of another adenoviral protein that inhibits p53 such as E4orf3 or E4orf6, could improve the outcome (50, 122). In line with these reports, we hypothesize that the introduction of SCM mutations into E2A, without the need for complete E2A deletion, could further improve the selectivity and efficacy of p53 selective oncolytic AdV.

Taken together, our findings unravel a novel mechanism by which HAdV inhibits p53-mediated transcriptional activation and apoptotic response of the host cell using HAdV DNA binding protein E2A. SUMOylated E2A promotes HAdV mediated inhibition of p53 dependent transcription by supporting p53 degradation and E1B-55K interaction with PML-V that leads to p53 SUMOylation and subsequent inhibition of the tumor suppressor activity.

## ACKNOWLEDGEMENT

We thank Ulrike Protzer, Thomas Floss, Martina Anton, Ruth Brack-Werner, Michelle Vincendeau and Percy Knolle for scientific discussions. Library preparation and Illumina RNAseq was performed at the RNAseq Core Facility, Institute of Human Genetics, Helmholtz Zentrum München. This work was funded by the Deutsche Forschungsgemeinschaft (DFG, German Research Foundation) in the framework of the Research Unit FOR5200 DEEP-DV (443644894) project SCHR 1479/5-1 and the Deutsche Krebshilfe e.V.. SS was funded by the Deutsche Forschungsgemeinschaft (DFG, German Research Foundation) under Germany’s Excellence Strategy - EXC 2155 - project number 390874280.

## AUTHOR CONTRIBUTION

Conceptualization: SS, MS; Methodology: MS, JM, VP, SH, LG, HCS, PG Investigation: MS, JM, VP, SH, LG, HCS, PG; Formal analysis: MS, HCS, SS; Resources: RTH, TD; Data Curation: MS, SS; Writing Original Draft: MS, SS; Writing – Reviewing & Editing: MS, SS; Visualization: MS, SS; Project administration: SS; Supervision: SS; Funding Acquisition: SS

## REFERENCES

1. Lane DP. Cancer. p53, guardian of the genome. Nature. 1992;358(6381):15–6.

2. Fischer M. Census and evaluation of p53 target genes. Oncogene. 2017;36(28):3943–56.

3. Horn HF, Vousden KH. Coping with stress: multiple ways to activate p53. Oncogene. 2007;26(9):1306–16.

4. Vousden KH, Prives C. Blinded by the Light: The Growing Complexity of p53. Cell. 2009;137(3):413–31.

5. Allen MA, Andrysik Z, Dengler VL, Mellert HS, Guarnieri A, Freeman JA, et al. Global analysis of p53-regulated transcription identifies its direct targets and unexpected regulatory mechanisms. Elife. 2014;3:e02200.

6. Sullivan KD, Galbraith MD, Andrysik Z, Espinosa JM. Mechanisms of transcriptional regulation by p53. Cell Death Differ. 2018;25(1):133–43.

7. Kress M, May E, Cassingena R, May P. Simian virus 40-transformed cells express new species of proteins precipitable by anti-simian virus 40 tumor serum. J Virol. 1979;31(2):472–83.

8. Lane DP, Crawford LV. T antigen is bound to a host protein in SV40-transformed cells. Nature. 1979;278(5701):261-3.

9. Linzer DI, Levine AJ. Characterization of a 54K dalton cellular SV40 tumor antigen present in SV40-transformed cells and uninfected embryonal carcinoma cells. Cell. 1979;17(1):43–52.

10. Smith AE, Smith R, Paucha E. Characterization of different tumor antigens present in cells transformed by simian virus 40. Cell. 1979;18(2):335–46.

11. Rivas C, Aaronson SA, Munoz-Fontela C. Dual Role of p53 in Innate Antiviral Immunity. Viruses. 2010;2(1):298–313.

12. Speir E, Modali R, Huang ES, Leon MB, Shawl F, Finkel T, et al. Potential role of human cytomegalovirus and p53 interaction in coronary restenosis. Science. 1994;265(5170):391-4.

13. Casavant NC, Luo MH, Rosenke K, Winegardner T, Zurawska A, Fortunato EA. Potential role for p53 in the permissive life cycle of human cytomegalovirus. J Virol. 2006;80(17):8390–401.

14. Hermannstädter A, Ziegler C, Kühl M, Deppert W, Tolstonog GV. Wild-type p53 enhances efficiency of simian virus 40 large-T-antigen-induced cellular transformation. J Virol. 2009;83(19):10106–18.

15. Izumi T, Io K, Matsui M, Shirakawa K, Shinohara M, Nagai Y, et al. HIV-1 viral infectivity factor interacts with TP53 to induce G2 cell cycle arrest and positively regulate viral replication. Proc Natl Acad Sci U S A. 2010;107(48):20798–803.

16. Liu X, Marmorstein R. When viral oncoprotein meets tumor suppressor: a structural view. Genes Dev. 2006;20(17):2332–7.

17. Maruzuru Y, Fujii H, Oyama M, Kozuka-Hata H, Kato A, Kawaguchi Y. Roles of p53 in herpes simplex virus 1 replication. J Virol. 2013;87(16):9323–32.

18. Zhu JY, Abate M, Rice PW, Cole CN. The ability of simian virus 40 large T antigen to immortalize primary mouse embryo fibroblasts cosegregates with its ability to bind to p53. J Virol. 1991;65(12):6872–80.

19. Ip WH, Dobner T. Cell transformation by the adenovirus oncogenes E1 and E4. FEBS Lett. 2020;594(12):1848–60.

20. Aloni-Grinstein R, Charni-Natan M, Solomon H, Rotter V. p53 and the Viral Connection: Back into the Future (‡). Cancers (Basel). 2018;10(6).

21. Trentin JJ, Yabe Y, Taylor G. The quest for human cancer viruses. Science. 1962;137(3533):835–41.

22. Kosulin K, Haberler C, Hainfellner JA, Amann G, Lang S, Lion T. Investigation of adenovirus occurrence in pediatric tumor entities. J Virol. 2007;81(14):7629–35.

23. Kosulin K, Hoffmann F, Clauditz TS, Wilczak W, Dobner T. Presence of adenovirus species C in infiltrating lymphocytes of human sarcoma. PLoS One. 2013;8(5):e63646.

24. Speiseder T, Hofmann-Sieber H, Rodríguez E, Schellenberg A, Akyüz N, Dierlamm J, et al. Efficient Transformation of Primary Human Mesenchymal Stromal Cells by Adenovirus Early Region 1 Oncogenes. J Virol. 2017;91(1).

25. Berciaud S, Rayne F, Kassab S, Jubert C, Faure-Della Corte M, Salin F, et al. Adenovirus infections in Bordeaux University Hospital 2008-2010: clinical and virological features. J Clin Virol. 2012;54(4):302–7.

26. Krilov LR. Adenovirus infections in the immunocompromised host. Pediatr Infect Dis J. 2005;24(6):555–6.

27. Lion T. Adenovirus infections in immunocompetent and immunocompromised patients. Clin Microbiol Rev. 2014;27(3):441–62.

28. Louie JK, Kajon AE, Holodniy M, Guardia-LaBar L, Lee B, Petru AM, et al. Severe pneumonia due to adenovirus serotype 14: a new respiratory threat? Clin Infect Dis. 2008;46(3):421–5.

29. Hofmann S, Mai J, Masser S, Groitl P, Herrmann A, Sternsdorf T, et al. ATO (Arsenic Trioxide) Effects on Promyelocytic Leukemia Nuclear Bodies Reveals Antiviral Intervention Capacity. Adv Sci (Weinh). 2020;7(8):1902130.

30. Lee CS, Bishop ES, Zhang R, Yu X, Farina EM, Yan S, et al. Adenovirus-Mediated Gene Delivery: Potential Applications for Gene and Cell-Based Therapies in the New Era of Personalized Medicine. Genes Dis. 2017;4(2):43–63.

31. St George JA. Gene therapy progress and prospects: adenoviral vectors. Gene Ther. 2003;10(14):1135–41.

32. Ganly I, Kirn D, Eckhardt G, Rodriguez GI, Soutar DS, Otto R, et al. A phase I study of Onyx-015, an E1B attenuated adenovirus, administered intratumorally to patients with recurrent head and neck cancer. Clin Cancer Res. 2000;6(3):798–806.

33. Goradel NH, Mohajel N, Malekshahi ZV, Jahangiri S, Najafi M, Farhood B, et al. Oncolytic adenovirus: A tool for cancer therapy in combination with other therapeutic approaches. J Cell Physiol. 2019;234(6):8636–46.

34. Ewer K, Sebastian S, Spencer AJ, Gilbert S, Hill AVS, Lambe T. Chimpanzee adenoviral vectors as vaccines for outbreak pathogens. Hum Vaccin Immunother. 2017;13(12):3020–32.

35. Tatsis N, Ertl HC. Adenoviruses as vaccine vectors. Mol Ther. 2004;10(4):616–29.

36. Ledgerwood JE, DeZure AD, Stanley DA, Coates EE, Novik L, Enama ME, et al. Chimpanzee Adenovirus Vector Ebola Vaccine. N Engl J Med. 2017;376(10):928–38.

37. Catanzaro AT, Koup RA, Roederer M, Bailer RT, Enama ME, Moodie Z, et al. Phase 1 safety and immunogenicity evaluation of a multiclade HIV-1 candidate vaccine delivered by a replication-defective recombinant adenovirus vector. J Infect Dis. 2006;194(12):1638–49.

38. Folegatti PM, Ewer KJ, Aley PK, Angus B, Becker S, Belij-Rammerstorfer S, et al. Safety and immunogenicity of the ChAdOx1 nCoV-19 vaccine against SARS-CoV-2: a preliminary report of a phase 1/2, single-blind, randomised controlled trial. Lancet. 2020;396(10249):467–78.

39. Hassan AO, Kafai NM, Dmitriev IP, Fox JM, Smith BK, Harvey IB, et al. A Single-Dose Intranasal ChAd Vaccine Protects Upper and Lower Respiratory Tracts against SARS-CoV-2. Cell. 2020;183(1):169–84.e13.

40. Mercado NB, Zahn R, Wegmann F, Loos C, Chandrashekar A, Yu J, et al. Single-shot Ad26 vaccine protects against SARS-CoV-2 in rhesus macaques. Nature. 2020;586(7830):583–8.

41. Bagchi S, Raychaudhuri P, Nevins JR. Adenovirus E1A proteins can dissociate heteromeric complexes involving the E2F transcription factor: a novel mechanism for E1A trans-activation. Cell. 1990;62(4):659–69.

42. Bandara LR, La Thangue NB. Adenovirus E1a prevents the retinoblastoma gene product from complexing with a cellular transcription factor. Nature. 1991;351(6326):494–7.

43. Blais A, Dynlacht BD. Hitting their targets: an emerging picture of E2F and cell cycle control. Curr Opin Genet Dev. 2004;14(5):527–32.

44. Ghosh MK, Harter ML. A viral mechanism for remodeling chromatin structure in G0 cells. Mol Cell. 2003;12(1):255–60.

45. Ait-Si-Ali S, Ramirez S, Barre FX, Dkhissi F, Magnaghi-Jaulin L, Girault JA, et al. Histone acetyltransferase activity of CBP is controlled by cycle-dependent kinases and oncoprotein E1A. Nature. 1998;396(6707):184–6.

46. Lowe SW, Ruley HE. Stabilization of the p53 tumor suppressor is induced by adenovirus 5 E1A and accompanies apoptosis. Genes Dev. 1993;7(4):535–45.

47. Nakajima T, Morita K, Tsunoda H, Imajoh-Ohmi S, Tanaka H, Yasuda H, et al. Stabilization of p53 by adenovirus E1A occurs through its amino-terminal region by modification of the ubiquitin-proteasome pathway. J Biol Chem. 1998;273(32):20036–45.

48. Zhang X, Turnell AS, Gorbea C, Mymryk JS, Gallimore PH, Grand RJ. The targeting of the proteasomal regulatory subunit S2 by adenovirus E1A causes inhibition of proteasomal activity and increased p53 expression. J Biol Chem. 2004;279(24):25122–33.

49. Dobner T, Horikoshi N, Rubenwolf S, Shenk T. Blockage by adenovirus E4orf6 of transcriptional activation by the p53 tumor suppressor. Science. 1996;272(5267):1470–3.

50. Soria C, Estermann FE, Espantman KC, O’Shea CC. Heterochromatin silencing of p53 target genes by a small viral protein. Nature. 2010;466(7310):1076–81.

51. Teodoro JG, Branton PE. Regulation of p53-dependent apoptosis, transcriptional repression, and cell transformation by phosphorylation of the 55-kilodalton E1B protein of human adenovirus type 5. J Virol. 1997;71(5):3620–7.

52. Yew PR, Berk AJ. Inhibition of p53 transactivation required for transformation by adenovirus early 1B protein. Nature. 1992;357(6373):82–5.

53. Yew PR, Liu X, Berk AJ. Adenovirus E1B oncoprotein tethers a transcriptional repression domain to p53. Genes Dev. 1994;8(2):190–202.

54. Endter C, Hartl B, Spruss T, Hauber J, Dobner T. Blockage of CRM1-dependent nuclear export of the adenovirus type 5 early region 1B 55-kDa protein augments oncogenic transformation of primary rat cells. Oncogene. 2005;24(1):55–64.

55. Endter C, Kzhyshkowska J, Stauber R, Dobner T. SUMO-1 modification required for transformation by adenovirus type 5 early region 1B 55-kDa oncoprotein. Proc Natl Acad Sci U S A. 2001;98(20):11312–7.

56. Cardoso FM, Kato SE, Huang W, Flint SJ, Gonzalez RA. An early function of the adenoviral E1B 55 kDa protein is required for the nuclear relocalization of the cellular p53 protein in adenovirus-infected normal human cells. Virology. 2008;378(2):339–46.

57. Liu Y, Colosimo AL, Yang XJ, Liao D. Adenovirus E1B 55-kilodalton oncoprotein inhibits p53 acetylation by PCAF. Mol Cell Biol. 2000;20(15):5540–53.

58. Pennella MA, Liu Y, Woo JL, Kim CA, Berk AJ. Adenovirus E1B 55-kilodalton protein is a p53-SUMO1 E3 ligase that represses p53 and stimulates its nuclear export through interactions with promyelocytic leukemia nuclear bodies. J Virol. 2010;84(23):12210–25.

59. Wimmer P, Berscheminski J, Blanchette P, Groitl P, Branton PE, Hay RT, et al. PML isoforms IV and V contribute to adenovirus-mediated oncogenic transformation by functionally inhibiting the tumor-suppressor p53. Oncogene. 2016;35(1):69–82.

60. Berscheminski J, Brun J, Speiseder T, Wimmer P, Ip WH, Terzic M, et al. Sp100A is a tumor suppressor that activates p53-dependent transcription and counteracts E1A/E1B-55K-mediated transformation. Oncogene. 2016;35(24):3178–89.

61. Blanchette P, Cheng CY, Yan Q, Ketner G, Ornelles DA, Dobner T, et al. Both BC-box motifs of adenovirus protein E4orf6 are required to efficiently assemble an E3 ligase complex that degrades p53. Mol Cell Biol. 2004;24(21):9619–29.

62. Querido E, Blanchette P, Yan Q, Kamura T, Morrison M, Boivin D, et al. Degradation of p53 by adenovirus E4orf6 and E1B55K proteins occurs via a novel mechanism involving a Cullin-containing complex. Genes Dev. 2001;15(23):3104–17.

63. Schreiner S, Wimmer P, Dobner T. Adenovirus degradation of cellular proteins. Future Microbiol. 2012;7(2):211–25.

64. Carter TH, Blanton RA. Possible role of the 72,000 dalton DNA-binding protein in regulation of adenovirus type 5 early gene expression. J Virol. 1978;25(2):664–74.

65. Babich A, Nevins JR. The stability of early adenovirus mRNA is controlled by the viral 72 kd DNA-binding protein. Cell. 1981;26(3 Pt 1):371–9.

66. Ahi YS, Vemula SV, Mittal SK. Adenoviral E2 IVa2 protein interacts with L4 33K protein and E2 DNA-binding protein. J Gen Virol. 2013;94(Pt 6):1325–34.

67. Nicolas JC, Sarnow P, Girard M, Levine AJ. Host range temperature-conditional mutants in the adenovirus DNA binding protein are defective in the assembly of infectious virus. Virology. 1983;126(1):228–39.

68. Puvion-Dutilleul F, Pedron J, Cajean-Feroldi C. Identification of intranuclear structures containing the 72K DNA-binding protein of human adenovirus type 5. Eur J Cell Biol. 1984;34(2):313–22.

69. Doucas V, Ishov AM, Romo A, Juguilon H, Weitzman MD, Evans RM, et al. Adenovirus replication is coupled with the dynamic properties of the PML nuclear structure. Genes Dev. 1996;10(2):196–207.

70. Everett RD. DNA viruses and viral proteins that interact with PML nuclear bodies. Oncogene. 2001;20(49):7266–73.

71. Ishov AM, Maul GG. The periphery of nuclear domain 10 (ND10) as site of DNA virus deposition. J Cell Biol. 1996;134(4):815–26.

72. Jul-Larsen A, Visted T, Karlsen BO, Rinaldo CH, Bjerkvig R, Lonning PE, et al. PML-nuclear bodies accumulate DNA in response to polyomavirus BK and simian virus 40 replication. Exp Cell Res. 2004;298(1):58–73.

73. Maul GG. Nuclear domain 10, the site of DNA virus transcription and replication. Bioessays. 1998;20(8):660–7.

74. Maul GG, Ishov AM, Everett RD. Nuclear domain 10 as preexisting potential replication start sites of herpes simplex virus type-1. Virology. 1996;217(1):67–75.

75. Carvalho T, Seeler JS, Ohman K, Jordan P, Pettersson U, Akusjärvi G, et al. Targeting of adenovirus E1A and E4-ORF3 proteins to nuclear matrix-associated PML bodies. J Cell Biol. 1995;131(1):45–56.

76. Hofmann S, Stubbe M, Mai J, Schreiner S. Double-edged Role of PML Nuclear Bodies during Human Adenovirus Infection. Virus Res. 2020:198280.

77. Stubbe M, Mai J, Paulus C, Stubbe HC, Berscheminski J, Karimi M, et al. Viral DNA Binding Protein SUMOylation Promotes PML Nuclear Body Localization Next to Viral Replication Centers. mBio. 2020;11(2).

78. Giard DJ, Aaronson SA, Todaro GJ, Arnstein P, Kersey JH, Dosik H, et al. In vitro cultivation of human tumors: establishment of cell lines derived from a series of solid tumors. J Natl Cancer Inst. 1973;51(5):1417–23.

79. Mitsudomi T, Oyama T, Gazdar AF, Minna JD, Okabayashi K, Shirakusa T. [Mutations of ras and p53 genes in human non-small cell lung cancer cell lines and their clinical significance]. Nihon Geka Gakkai zasshi. 1992;93(9):944–7.

80. Catalucci D, Sporeno E, Cirillo A, Ciliberto G, Nicosia A, Colloca S. An adenovirus type 5 (Ad5) amplicon-based packaging cell line for production of high-capacity helper-independent deltaE1-E2-E3-E4 Ad5 vectors. J Virol. 2005;79(10):6400–9.

81. Tatham MH, Rodriguez MS, Xirodimas DP, Hay RT. Detection of protein SUMOylation in vivo. Nat Protoc. 2009;4(9):1363–71.

82. Schreiner S, Wimmer P, Sirma H, Everett RD, Blanchette P, Groitl P, et al. Proteasome-dependent degradation of Daxx by the viral E1B-55K protein in human adenovirus-infected cells. J Virol. 2010;84(14):7029–38.

83. Groitl P, Dobner T. Construction of adenovirus type 5 early region 1 and 4 virus mutants. Methods Mol Med. 2007;130:29–39.

84. Kindsmuller K, Groitl P, Hartl B, Blanchette P, Hauber J, Dobner T. Intranuclear targeting and nuclear export of the adenovirus E1B-55K protein are regulated by SUMO1 conjugation. Proceedings of the National Academy of Sciences of the United States of America. 2007;104(16):6684–9.

85. Kindsmuller K, Schreiner S, Leinenkugel F, Groitl P, Kremmer E, Dobner T. A 49-kilodalton isoform of the adenovirus type 5 early region 1B 55-kilodalton protein is sufficient to support virus replication. J Virol. 2009;83(18):9045–56.

86. Wimmer P, Schreiner S, Everett RD, Sirma H, Groitl P, Dobner T. SUMO modification of E1B-55K oncoprotein regulates isoform-specific binding to the tumour suppressor protein PML. Oncogene. 2010;29(40):5511–22.

87. Berscheminski J, Wimmer P, Brun J, Ip WH, Groitl P, Horlacher T, et al. Sp100 isoform-specific regulation of human adenovirus 5 gene expression. J Virol. 2014;88(11):6076–92.

88. Langmead B, Trapnell C, Pop M, Salzberg SL. Ultrafast and memory-efficient alignment of short DNA sequences to the human genome. Genome Biol. 2009;10(3):R25.

89. Zhang Y, Liu T, Meyer CA, Eeckhoute J, Johnson DS, Bernstein BE, et al. Model-based analysis of ChIP-Seq (MACS). Genome Biol. 2008;9(9):R137.

90. Quinlan AR, Hall IM. BEDTools: a flexible suite of utilities for comparing genomic features. Bioinformatics. 2010;26(6):841–2.

91. Heinz S, Benner C, Spann N, Bertolino E, Lin YC, Laslo P, et al. Simple combinations of lineage-determining transcription factors prime cis-regulatory elements required for macrophage and B cell identities. Mol Cell. 2010;38(4):576–89.

92. Shen L, Shao NY, Liu X, Maze I, Feng J, Nestler EJ. diffReps: detecting differential chromatin modification sites from ChIP-seq data with biological replicates. PLoS One. 2013;8(6):e65598.

93. Lerdrup M, Johansen JV, Agrawal-Singh S, Hansen K. An interactive environment for agile analysis and visualization of ChIP-sequencing data. Nat Struct Mol Biol. 2016;23(4):349–57.

94. Machanick P, Bailey TL. MEME-ChIP: motif analysis of large DNA datasets. Bioinformatics. 2011;27(12):1696–7.

95. Haack TB, Kopajtich R, Freisinger P, Wieland T, Rorbach J, Nicholls TJ, et al. ELAC2 mutations cause a mitochondrial RNA processing defect associated with hypertrophic cardiomyopathy. Am J Hum Genet. 2013;93(2):211–23.

96. Bray NL, Pimentel H, Melsted P, Pachter L. Near-optimal probabilistic RNA-seq quantification. Nat Biotechnol. 2016;34(5):525–7.

97. Yates AD, Achuthan P, Akanni W, Allen J, Allen J, Alvarez-Jarreta J, et al. Ensembl 2020. Nucleic Acids Res. 2020;48(D1):D682–d8.

98. Love MI, Huber W, Anders S. Moderated estimation of fold change and dispersion for RNA-seq data with DESeq2. Genome Biol. 2014;15(12):550.

99. Bosari S, Viale G, Roncalli M, Graziani D, Borsani G, Lee AK, et al. p53 gene mutations, p53 protein accumulation and compartmentalization in colorectal adenocarcinoma. Am J Pathol. 1995;147(3):790–8.

100. Moll UM, LaQuaglia M, Benard J, Riou G. Wild-type p53 protein undergoes cytoplasmic sequestration in undifferentiated neuroblastomas but not in differentiated tumors. Proc Natl Acad Sci U S A. 1995;92(10):4407–11.

101. Moll UM, Riou G, Levine AJ. Two distinct mechanisms alter p53 in breast cancer: mutation and nuclear exclusion. Proc Natl Acad Sci U S A. 1992;89(15):7262–6.

102. Ornelles DA, Shenk T. Localization of the adenovirus early region 1B 55-kilodalton protein during lytic infection: association with nuclear viral inclusions requires the early region 4 34-kilodalton protein. J Virol. 1991;65(1):424–9.

103. Hendriks IA, Vertegaal AC. A comprehensive compilation of SUMO proteomics. Nat Rev Mol Cell Biol. 2016;17(9):581–95.

104. Leppard KN, Everett RD. The adenovirus type 5 E1b 55K and E4 Orf3 proteins associate in infected cells and affect ND10 components. J Gen Virol. 1999;80 (Pt 4):997–1008.

105. von Stromberg K, Seddar L, Ip WH, Günther T, Gornott B, Weinert SC, et al. The human adenovirus E1B-55K oncoprotein coordinates cell transformation through regulation of DNA-bound host transcription factors. Proc Natl Acad Sci U S A. 2023;120(44):e2310770120.

106. Zhao H, Chen M, Valdés A, Lind SB, Pettersson U. Transcriptomic and proteomic analyses reveal new insights into the regulation of immune pathways during adenovirus type 2 infection. BMC Microbiol. 2019;19(1):15.

107. Zhao H, Dahlö M, Isaksson A, Syvänen AC, Pettersson U. The transcriptome of the adenovirus infected cell. Virology. 2012;424(2):115–28.

108. Aubrey BJ, Kelly GL, Janic A, Herold MJ, Strasser A. How does p53 induce apoptosis and how does this relate to p53-mediated tumour suppression? Cell Death Differ. 2018;25(1):104–13.

109. Kanehisa M, Furumichi M, Sato Y, Matsuura Y, Ishiguro-Watanabe M. KEGG: biological systems database as a model of the real world. Nucleic Acids Res. 2025;53(D1):D672–d7.

110. Kanehisa M. Toward understanding the origin and evolution of cellular organisms. Protein Sci. 2019;28(11):1947–51.

111. Kanehisa M, Goto S. KEGG: kyoto encyclopedia of genes and genomes. Nucleic Acids Res. 2000;28(1):27–30.

112. White E. Mechanisms of apoptosis regulation by viral oncogenes in infection and tumorigenesis. Cell Death Differ. 2006;13(8):1371–7.

113. Sato N, Uematsu M, Fujimoto K, Uematsu S, Imoto S. ggkegg: analysis and visualization of KEGG data utilizing the grammar of graphics. Bioinformatics. 2023;39(10).

114. Wang B, Lam TH, Soh MK, Ye Z, Chen J, Ren EC. Influenza A Virus Facilitates Its Infectivity by Activating p53 to Inhibit the Expression of Interferon-Induced Transmembrane Proteins. Front Immunol. 2018;9:1193.

115. Whyte P, Buchkovich KJ, Horowitz JM, Friend SH, Raybuck M, Weinberg RA, et al. Association between an oncogene and an anti-oncogene: the adenovirus E1A proteins bind to the retinoblastoma gene product. Nature. 1988;334(6178):124–9.

116. DeCaprio JA, Ludlow JW, Figge J, Shew JY, Huang CM, Lee WH, et al. SV40 large tumor antigen forms a specific complex with the product of the retinoblastoma susceptibility gene. Cell. 1988;54(2):275–83.

117. Sarnow P, Ho YS, Williams J, Levine AJ. Adenovirus E1b-58kd tumor antigen and SV40 large tumor antigen are physically associated with the same 54 kd cellular protein in transformed cells. Cell. 1982;28(2):387–94.

118. Talotta F, Casalino L, Verde P. The nuclear oncoprotein Fra-1: a transcription factor knocking on therapeutic applications’ door. Oncogene. 2020;39(23):4491–506.

119. Sakurai F, Tachibana M, Mizuguchi H. Adenovirus vector-based vaccine for infectious diseases. Drug Metab Pharmacokinet. 2022;42:100432.

120. Abudoureyimu M, Lai Y, Tian C, Wang T, Wang R, Chu X. Oncolytic Adenovirus—A Nova for Gene-Targeted Oncolytic Viral Therapy in HCC. Frontiers in Oncology. 2019;9(1182).

121. Bischoff JR, Kirn DH, Williams A, Heise C, Horn S, Muna M, et al. An adenovirus mutant that replicates selectively in p53-deficient human tumor cells. Science. 1996;274(5286):373–6.

122. Yanagawa-Matsuda A, Mikawa Y, Habiba U, Kitamura T, Yasuda M, Towfik-Alam M, et al. Oncolytic potential of an E4-deficient adenovirus that can recognize the stabilization of AU-rich element containing mRNA in cancer cells. Oncol Rep. 2019;41(2):954–60.

