## Supplementary figures and images for "Human Adenovirus (HAdV) mediated inhibition of the tumor suppressor p53 is controlled by SUMOylation of the viral E2A/DBP protein"

### Suppl Fig. 1

#### Supplement Figure 1
Stubbe et al.

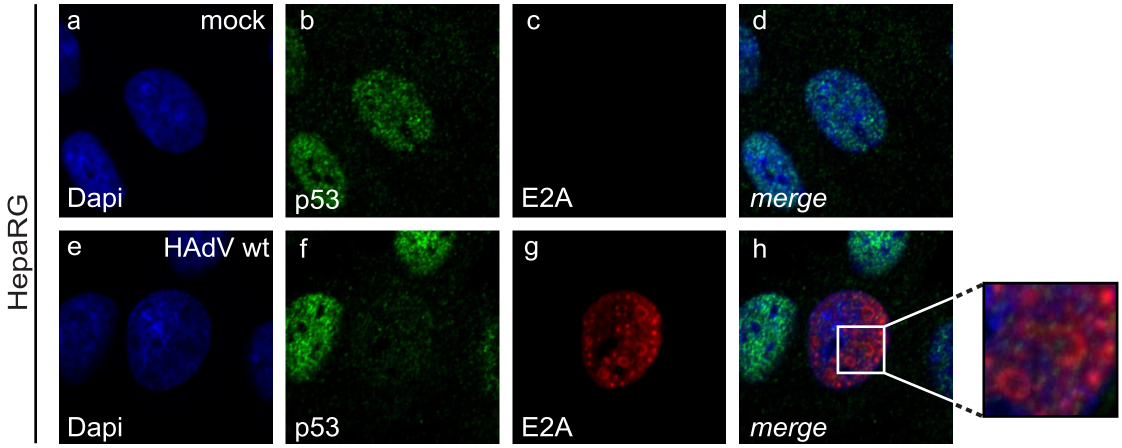

### Suppl Fig. 2

Supplement Figure 2
Stubbe et al.

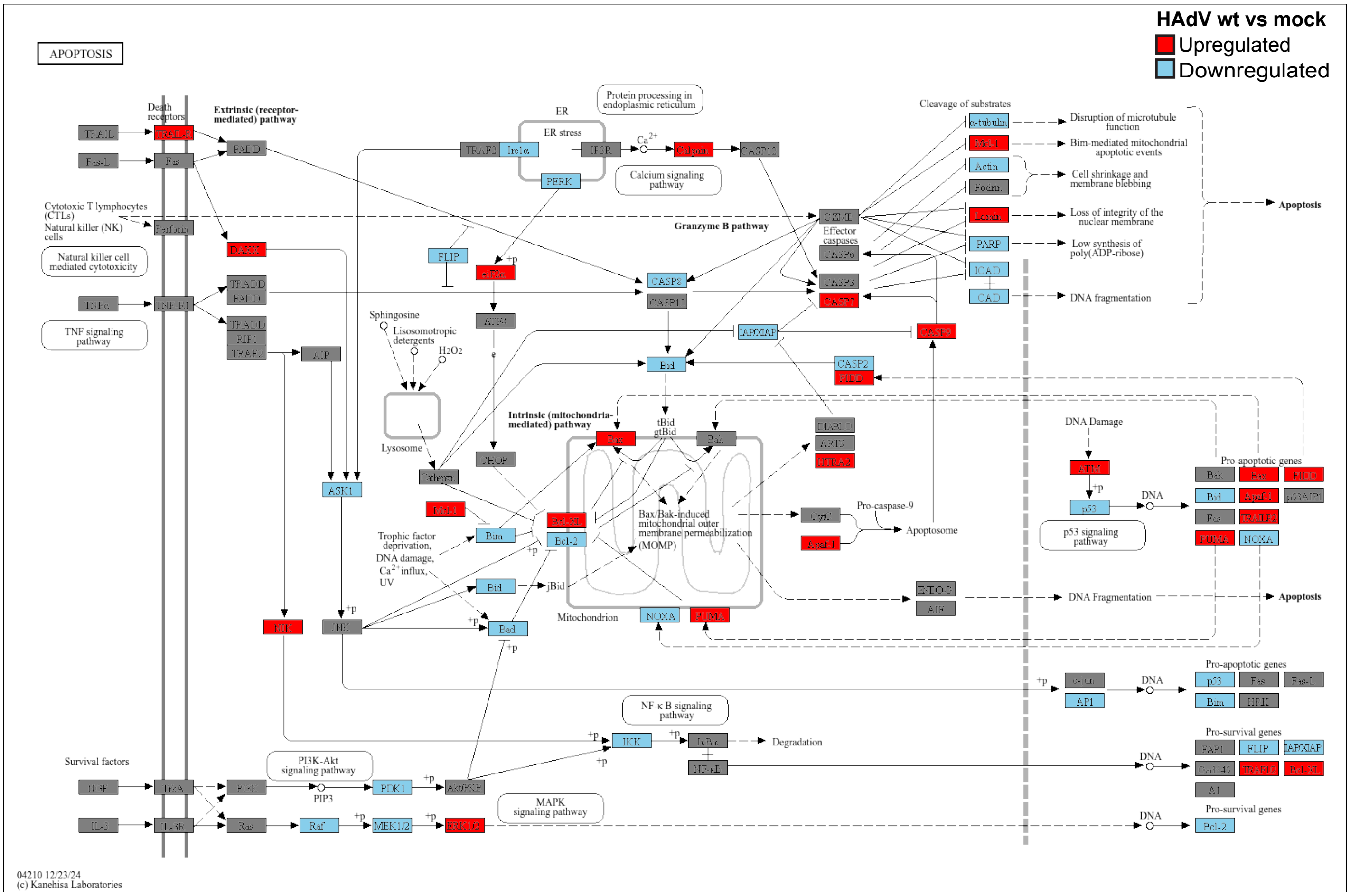
