## Supplementary material for "Human Adenovirus (HAdV) mediated inhibition of the tumor suppressor p53 is controlled by SUMOylation of the viral E2A/DBP protein": Suppl Figure legends

### SUPPLEMENT FIGURE LEGENDS

**Suppl. Fig. 1. p53 does not colocalize with the viral RCs 36 h post HAdV infection.** HepaRG cells were infected with HAdV wt virus at a multiplicity of infection of 20 FFU/cell. The cells were fixed with 2% PFA 36h p.i. and double labeled with mAb rb (α-E2A) and mAb DO-1 (α-p53). Primary Abs were detected with Alexa647 (E2A, red) and Alexa488 (α-p53, green) conjugated secondary Abs. Nuclear staining was performed using Dapi. Anti-p53 (green; panels b, f) and anti-E2A (red; panels c, g), staining patterns representative of at least 50 analyzed cells are shown. Overlays of single images (merge) are shown in panels d and h.

**Suppl. Fig. 2. HAdV mediated manipulation of apoptosis pathway.** Data obtained by transcriptomic analysis of differentially expressed transcripts in HAdV wt infected cells compared to mock infected cells was rendered on KEGG apoptosis pathway graph using ggKEGG (113). Most significantly expressed transcript of each gene (FDR $< 0.1$) was rendered on the KEGG apoptosis pathway graph. Red color signifies upregulation, blue downregulation.
